# Microbial Tryptophan Metabolism Activates Host Lysosomal Activity to Facilitate Lipid Breakdown and Ameliorate Hepatic Steatosis

**DOI:** 10.1101/2025.06.28.662086

**Authors:** Kenan Zhang, Zihan Luo, Yan Li, Yan Chen, Lang Wang, Yanan Liu, Ruizhi Yang, Qian Li, Jiahao Zhao, Bin Qi, Zhao Shan

## Abstract

Lysosomes are central to lipid metabolism, yet how gut microbiota-derived metabolites regulate lysosomal function to influence host lipid homeostasis remains unknown. Here, we identify an evolutionarily conserved mechanism in which bacterial tryptophan metabolism activates lysosomal activity to promote lipid breakdown. By developing a lysosomal-responsive lipid reporter in *C. elegans* to screen for bacterial metabolic states that modulate host lipid storage, we discover that *E. coli* tryptophan catabolism via tryptophanase TnaA induces lysosomal lipid chaperone LBP-8, driving lipid mobilization. Moreover, tryptophan metabolites enhanced lysosomal acidification and degradation capacity, while genetic disruption of lysosomal regulators reversed these effects. Strikingly, bacterial tryptophan metabolism further promoted mitochondrial β-oxidation through lysosomal lipase activity. This pathway was conserved in mammalian hepatocytes, where *E. coli*-derived tryptophan metabolites enhance lysosomal function and reduce lipid accumulation. In high-fat diet mice, restoring gut bacterial tryptophan metabolism alleviated hepatic steatosis. Our work uncovers microbiota-regulated lysosomal activation as a critical axis in lipid homeostasis, highlighting its potential as a therapeutic target for metabolic disorders linked to lysosomal dysfunction.

**Significance:** We uncover a conserved mechanism by which microbial tryptophan metabolism enhances lysosomal function to maintain host lipid homeostasis. Specifically, we demonstrate that bacterial tryptophan catabolism—via the enzyme TnaA—promotes lysosomal acidification, proteolytic capacity, and structural remodeling in *C. elegans*, driving lipid breakdown through the lysosomal chaperone LBP-8. This activation boosts mitochondrial β-oxidation and reduces lipid storage. Importantly, the same pathway operates in mammalian hepatocytes and in a high-fat diet mouse model, where restoring bacterial tryptophan metabolism markedly alleviates hepatic steatosis. Our findings bridge microbial metabolism and lysosomal dynamics, offering fresh insights into host–microbe crosstalk and metabolic regulation.

**Highlights:** TnaA-mediated bacterial tryptophan catabolism promotes lipid mobilization via lysosomal chaperone.

Bacterial tryptophan metabolites boost lysosomal function, lipid breakdown, and mitochondrial β-oxidation.

Conserved microbiota-lysosome-lipid axis from worms to mammalian liver.

Restoring gut bacterial tryptophan metabolism alleviates hepatic steatosis in high-fat diet mice.

## Introduction

The gut microbiota plays a pivotal role in regulating host metabolism, including lipid homeostasis, through the production of bioactive metabolites that influence cellular processes such as energy storage, mitochondrial function, and organelle dynamics (Krautkramer et al., 2021; Lee et al., 2024). Among these processes, lysosomes serve as central hubs for lipid degradation, where acid hydrolases break down complex lipids into free fatty acids for energy production or recycling (Settembre and Ballabio, 2014b). Mitochondria, essential for energy production and evolutionarily derived from bacteria (Roger et al., 2017), engage in critical communication with bacteria that regulates host immune responses (Erny et al., 2021), metabolism (Qi and Han, 2018; Tian and Han, 2022) and aging (Han et al., 2017). While the impact of bacterial metabolites on mitochondria is becoming increasingly in recent years (Lee et al., 2024), the specific regulatory roles of these metabolites on lysosomal function remain largely unexplored.

Lysosomes are central to lipid metabolism, acting as enzymatic hubs for fat breakdown and recycling. In mammals, lysosomal acid lipase (LAL) is indispensable, degrading triglycerides (TGs), cholesterol esters (CEs), and lipoproteins in the acidic lysosomal environment (Settembre and Ballabio, 2014a). In *C. elegans,* major fats are stored in vesicles distinct from lysosome-related organelles (O’Rourke et al., 2009). Lysosomal lipases are essential for lipid droplet (LD) degradation, as their absence results in LD accumulation (O’Rourke and Ruvkun, 2013). Beyond breakdown, these lipases regulate lipid signaling: LIPL-4 activation triggers a lysosome-to-nucleus cascade that boosts lipolysis and extends lifespan (Folick et al., 2015). This pathway involves LIPL-4 and its lipid chaperone LBP-8, which mobilizes fatty acids to enhance mitochondrial β-oxidation, reduce lipid storage, and promote longevity (Ramachandran et al., 2019). Dysregulation of lysosomal function disrupts lipid homeostasis and is implicated in pathologies ranging from lysosomal storage disorders (Platt et al., 2012) to metabolic diseases (Gros and Muller, 2023). Despite their central role in lipid handling, the mechanisms by which extrinsic factors—such as gut microbial metabolites—modulate lysosomal activity remain poorly understood. Identifying bacterial metabolites that fine-tune lysosomal function is thus essential for unraveling host-microbe metabolic crosstalk and developing therapies for lysosomal-related disorders.

Gut bacterial metabolism of tryptophan plays a pivotal role in animal physiology, generating indole derivatives and other metabolites that regulate immune responses, intestinal barrier integrity, and systemic metabolism (Agus et al., 2018; Hezaveh et al., 2022; Scott et al., 2020; Tintelnot et al., 2023). In metabolic disorders such as obesity, type 2 diabetes (T2D), and non-alcoholic fatty liver disease (NAFLD), compositional and functional alterations in the gut microbiota are well-documented (Sonnenburg and Bäckhed, 2016), with impaired bacterial tryptophan metabolism emerging as a key contributor to disease pathogenesis (Natividad et al., 2018). However, how microbial tryptophan metabolism regulates cellular organelle dynamics in host is still unclear. Given the central role of lysosomes in lipid catabolism and their dysregulation in metabolic diseases, understanding whether gut-derived tryptophan metabolites modulate lysosomal activity could unveil novel therapeutic targets. Addressing this gap is critical for leveraging host-microbe crosstalk to restore lipid metabolism and combat lysosomal-related disorders.

In this study, we developed a lysosomal-responsive lipid reporter system in *C. elegans* to screen for bacterial metabolic states that modulate host lipid storage. Our findings reveal an evolutionarily conserved mechanism in which bacterial tryptophan metabolism activates lysosomal activity to drive lipid catabolism—a process validated in both *C. elegans* and mammalian models. These results deepen our understanding of inter-organismal metabolic crosstalk and establish a framework for therapies targeting microbial tryptophan metabolic pathways to regulate lysosomal function and counteract lipid-related pathologies.

## Results

### Establishment of a Lysosomal-responsive Lipid Reporter System to Screen Bacterial Metabolic State Changes Regulating Lipid Metabolism

The constitutive activation of lysosomal lipolysis via overexpression of lysosomal acid lipase LIPL-4 drives nuclear translocation of signaling molecules, including the lipid chaperone LBP-8 and the lipid messenger oleoylethanolamine, which transcriptionally activate metabolic genes to promote fat mobilization (Folick et al., 2015). Specifically, the LIPL-4–LBP-8 lysosomal signaling pathway enhances mitochondrial β-oxidation, leading to reduced total fat storage in animals overexpressing *lipl-4* or its downstream effector *lbp-8* (Ramachandran et al., 2019). This suggests an inverse correlation between *lbp-8* expression levels and lipid accumulation, where elevated *lbp-8* expression correlates with reduced lipid stores (Figure 1A). To validate this relationship, we generated transgenic *C. elegans* overexpressing *lbp-8* and quantified lipid levels using Oil Red O (ORO) staining. As predicted, *lbp-8*-overexpressing animals (*lbp-8 Tg*) exhibited significantly reduced lipid accumulation compared to wild-type controls (Figure 1B).

**Figure 1.**
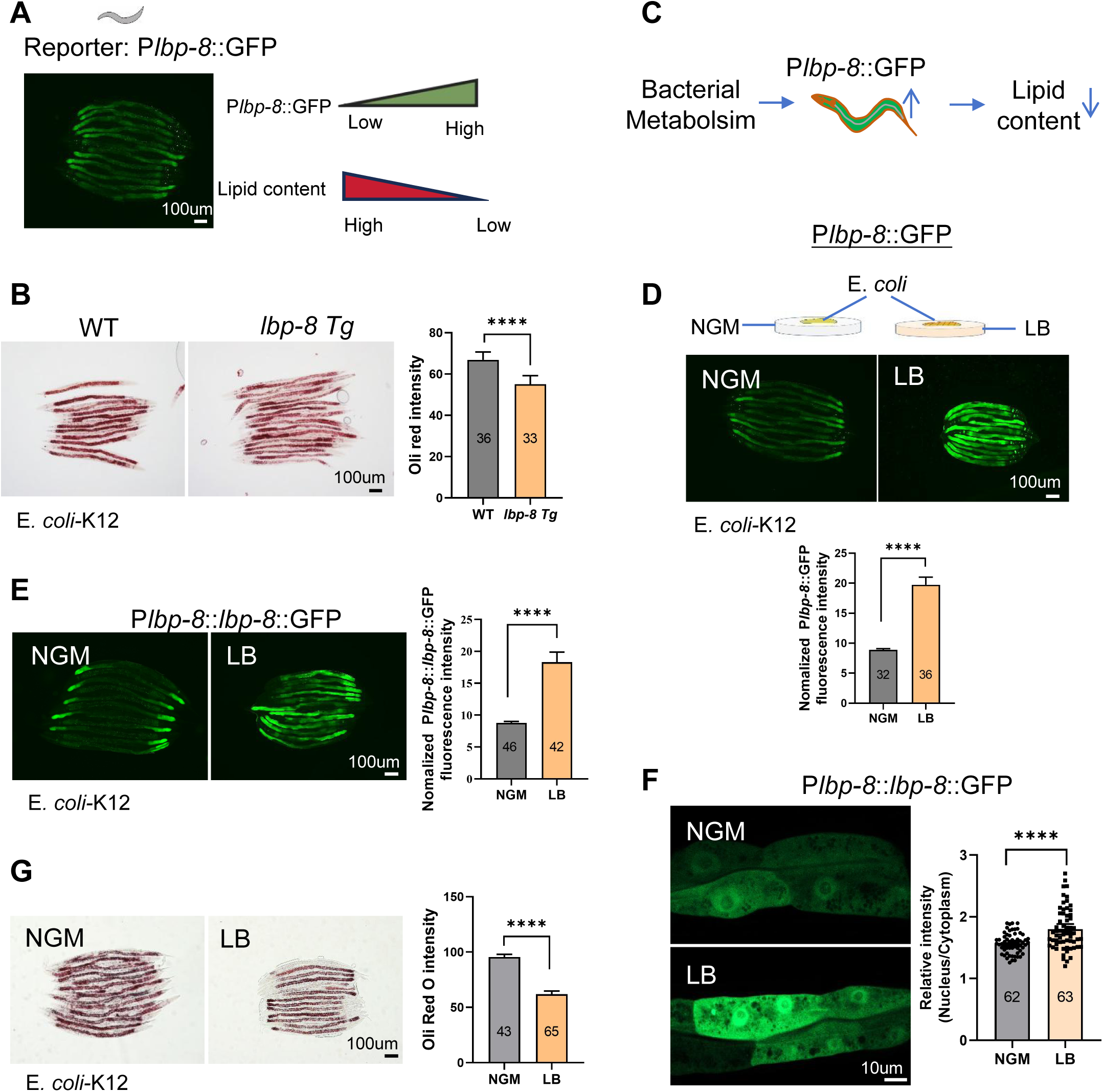
Establishment of a lysosomal-responsive lipid reporter in *C. elegans* to screen bacterial metabolic state changes regulating lipid metabolism. **(A)** Schematic of the inverse relationship between *lbp-8* expression and fat accumulation, alongside representative fluorescence micrographs of the P*lbp-8*::GFP reporter in wild-type L4 larvae. Scale bar, 100 µm. **(B)** Oil Red O staining and quantification of total lipid levels in wild-type (WT) versus *lbp-8*–overexpressing animals (*lbp-8 Tg*) at the L4 stage. The number of animals analyzed is indicated. Scale bar, 100 µm. **(C)** Schematic model illustrating that changes in the bacterial metabolic state induce *lbp-8* expression, thereby promoting lipid breakdown. **(D)** Fluorescence micrographs and quantification of P*lbp-8*::GFP reporter expression in L4 larvae fed on standard NGM plates seeded with *E. coli* K12 (NGM) versus LB-conditioned *E. coli* K12 (LB). The number of animals analyzed is indicated. Scale bar, 100 µm. **(E)** Fluorescence images and quantification of the P*lbp-8*::*lbp-8*::GFP expression in L4 larvae under the same dietary conditions as in (D). The number of animals analyzed is indicated. Scale bar, 100 µm. **(F)** High-magnification fluorescence images and quantification of the nuclear-to-cytoplasmic GFP intensity ratio for LBP-8::GFP in L4 larvae on NGM versus LB *E. coli*. The number of animals analyzed is indicated. Scale bar, 10 µm. **(G)** Oil Red O staining and quantification of lipid content in wild-type L4 larvae fed on NGM versus LB *E. coli-*K12. The number of animals analyzed is indicated.Scale bar, 100 µm. Data represent mean ± SD. All statistical analyses were performed using unpaired two-tailed Student’s t-test. ****p < 0.0001. All experiments were performed independently at least three times. See also Figure S1.

To determine whether bacterial metabolic states influence *lbp-8* expression and subsequent lipid metabolism, we employed an reporter which involved in LIPL-4–LBP-8 lysosomal signaling pathway for lipid metabolism (Ramachandran et al., 2019), P*lbp-8*::GFP transcriptional reporter (Figure 1C). We first tested this by culturing *C. elegans* under standard conditions on NGM plates seeded with metabolically compromised *E. coli* (either antibiotic-treated with ampicillin or UV-killed). However, no significant changes in P*lbp-8*::GFP expression were observed under these conditions (Figure S1A).

Remarkably, when we cultured worms on nutrient-rich LB medium plates seeded with *E. coli*-K12, we observed significant induction of *lbp-8* at both the transcriptional (Figure 1D) and translational (Figure 1E) levels. Consistent with the role of LBP-8 in lysosomal signaling, nuclear translocation of LBP-8—a hallmark of lysosomal lipolysis activation (Folick et al., 2015)—was significantly enhanced in animals fed *E. coli* on LB plates (Figure 1F). Correspondingly, Oil Red O (ORO) staining revealed significantly reduced lipid stores in these animals (Figure 1G).

The LB medium condition promoted robust bacterial overgrowth, resulting in thick bacterial lawns (Figure S1B), suggesting that the nutrient-rich environment may alter *E. coli* metabolism that activate host lipid catabolism. Together, these findings suggested that bacterial metabolic changes induced by LB culture conditions can trigger LBP-8 expression and promote lipid breakdown in *C. elegans*.

### Bacterial Tryptophan Metabolism Regulates Host Lipid Metabolism via Induction of LBP-8

To identify bacterial metabolic factors modulating *lbp-8* activation and lipid mobilization in *C. elegans*, we conducted a genome-wide screen of the *E. coli* single-gene knockout library (Keio collection). Using the parental strain *E. coli* K-12 BW25113 as a control, we assessed P*lbp-8::*GFP expression under LB-medium plate culture (Figure 2A). After screening an *E. coli* single-gene knockout library, we found that 19 *E. coli* mutant significantly reduced *lbp-8* expression (Figure 2A, Figuer S2A).

**Figure 2.**
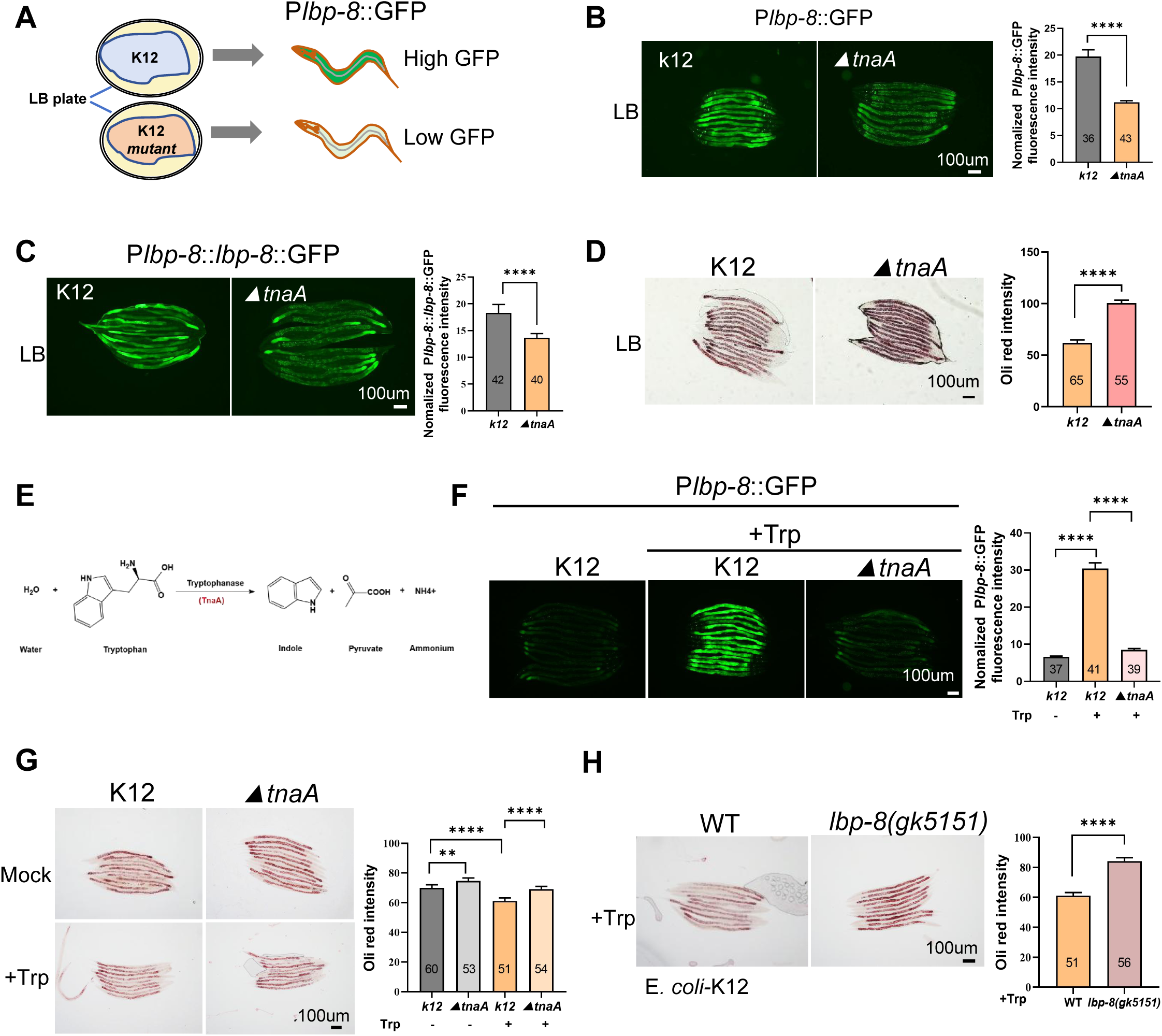
Bacterial tryptophan metabolism regulates lipid homeostasis. **(A)** Schematic of a genome-wide screen using the *E. coli* single-gene knockout library in *C. elegans* harboring the P*lbp-8*::GFP reporter, identifying *E. coli* mutants that fail to induce GFP on LB medium. **(B)** Representative fluorescence images and quantification of the P*lbp-8*::GFP reporter in L4 animals feed on LB medium seeded with wild-type *E. coli* K12 or the Δ*tnaA* mutant. The number of animals analyzed is indicated. Scale bar, 100 μm. **(C)** Fluorescence images and quantification of the P*lbp-8*::*lbp-8*::GFP reporter under the same conditions as in (B). The number of animals analyzed is indicated. Scale bar, 100 μm. **(D)** Oil Red O staining and quantification of lipid content in wild-type L4 animals feed on LB medium with wild-type *E. coli* K12 or Δ*tnaA*. The number of animals analyzed is indicated. Scale bar, 100 μm. **(E)** Diagram of the tryptophan catabolic pathway in *E. coli*, highlighting the role of TnaA. **(F)** Fluorescence images and quantification of the P*lbp-8*::GFP reporter in L4 animals feed on NGM plate seeded with *E. coli*-K12, or 10 mM tryptophan supplemented NGM plated seeded with *E. coli*-K12 or Δ*tnaA*. The number of animals analyzed is indicated. Scale bar, 100 μm. **(G)** Oil Red O staining and quantification of lipid levels in wild-type L4 larvae fed on NGM plate with or without 10 mM tryptophan, seeded with K12 or Δ*tnaA*.The number of animals analyzed is indicated. Scale bar, 100 μm. **(H)** Oil Red O staining and quantification of wild-type and *lbp-8*(gk5151) mutant L4 worms fed on NGM plate supplemented with 10 mM tryptophan and seeded with K12. The number of animals analyzed is indicated. Scale bar, 100 μm. Data represent mean ± SD. All statistical analyses were performed using unpaired two-tailed Student’s t-test. ****p < 0.0001, **p < 0.01. All experiments were performed independently at least three times. See also Figure S2.

The *E. col*-tnaA mutant exhibited the robust effect on *lbp-8* expression. Deletion of tnaA markedly suppressed *lbp-8* expression at both transcriptional (Figure 2B) and translational (Figure 2C) levels under LB-medium plate conditions. Transcriptomic analysis confirmed that tnaA deficiency abolished LB-induced *lbp-8* upregulation (Figures S2B), while nuclear translocation of LBP-8 was similarly reduced in animals fed *E. coli*-tnaA (Figure S2C). Notably, Oil Red O (ORO) staining revealed elevated total lipid levels in *C. elegans* fed tnaA mutants on LB plates (Figure 2D), indicating that *E. coli* TnaA-dependent metabolism promotes *lbp-8* expression and subsequent lipid breakdown.

TnaA, a tryptophanase, catalyzes tryptophan conversion into indole, pyruvate, and ammonium (Figure 2E). Since LB plates are rich in tryptone—a source of amino acids including tryptophan—this raised the possibility that bacterial tryptophan metabolism modulates host lipid breakdown by inducing *lbp-8* expression. To test this, we supplemented standard NGM plates (seeded with wild-type *E. coli*) with tryptophan, which robustly induced *lbp-8* expression (Figure 2F, Figures S2D, S2E) and LBP-8 nuclear translocation (Figure S2F). In contrast, supplementing tryptophan into plates with heat-killed *E. coli* did not induce *lbp-8* expression (Figure S2G), underscoring the necessity of active bacterial metabolism for *lbp-8* regulation.

Furthermore, while tryptophan supplementation induced *lbp-8* expression in animals fed wild-type *E. coli*, this induction was lost when worms were fed the *E. coli* -tnaA mutant (Figure 2F). Correspondingly, lipid levels decreased in wild-type-fed worms with tryptophan supplementation (Figure 2G) but were restored in animals fed the tnaA mutant. Finally, *lbp-8* mutant animals exhibited higher lipid levels despite tryptophan supplementation when fed wild-type *E. coli* (Figure 2H), suggesting that bacterial tryptophan metabolism promotes lipid breakdown via LBP-8-dependent pathway.

In summary, our results demonstrate that bacterial TnaA-mediated tryptophan metabolism induces *lbp-8* expression, which in turn drives lipid breakdown in *C. elegans*.

### Bacterial Tryptophan Metabolism Induce Lysosomal-related Genes Expression

To further elucidate the molecular mechanisms by which bacterial tryptophan metabolism modulates host lipid metabolism, we performed RNA-sequencing (RNA-seq) on *C. elegans* under two experimental conditions: (1) wild-type animals fed *E. coli* K12 (Mock-K12) versus those fed K12 supplemented with tryptophan (Trp-K12), and (2) Trp-K12 versus animals fed tnaA mutants in tryptophan-supplemented medium (Trp-tnaA) (Figure S3A). The transcriptomic analysis revealed that tryptophan supplementation in the K12 group upregulated approximately 2,050 genes while downregulating 1,285 genes compared to the mock treatment. In contrast, comparing Trp-K12 to Trp-tnaA animals showed 1648 genes upregulated and 1437 downregulated (Figure S3B, Table S1), indicating that the presence of bacterial TnaA is essential for a full transcriptional response to tryptophan.

Focusing on lipid metabolism, we observed that key lipases responsible for the breakdown of triacylglycerol into free fatty acids—including adipose triglyceride lipase (ATGL/*atgl-1*) and hormone-sensitive lipase (HSL/*hosl-1*)—as well as lysosomal acid lipases, were significantly induced by tryptophan metabolism (Figure 3A). Additionally, the lipid-binding proteins LBP-1 and LBP-8 were specifically induced, further linking bacterial tryptophan metabolism to enhanced lipid mobilization. In contrast, genes encoding enzymes involved in lipid synthesis, such as POD-2 (acetyl-CoA carboxylase), FASN-1 (fatty acid synthase), several elongases (*elo-2*, *elo-8*), and desaturases (*fat-6*, *fat-7*), remained unchanged or were suppressed (Figure 3A). This result suggested that bacterial tryptophan metabolic mainly induce lysosomal lipases expression which may facilitate breakdown of triacylglycerol into free fatty acids.

**Figure 3.**
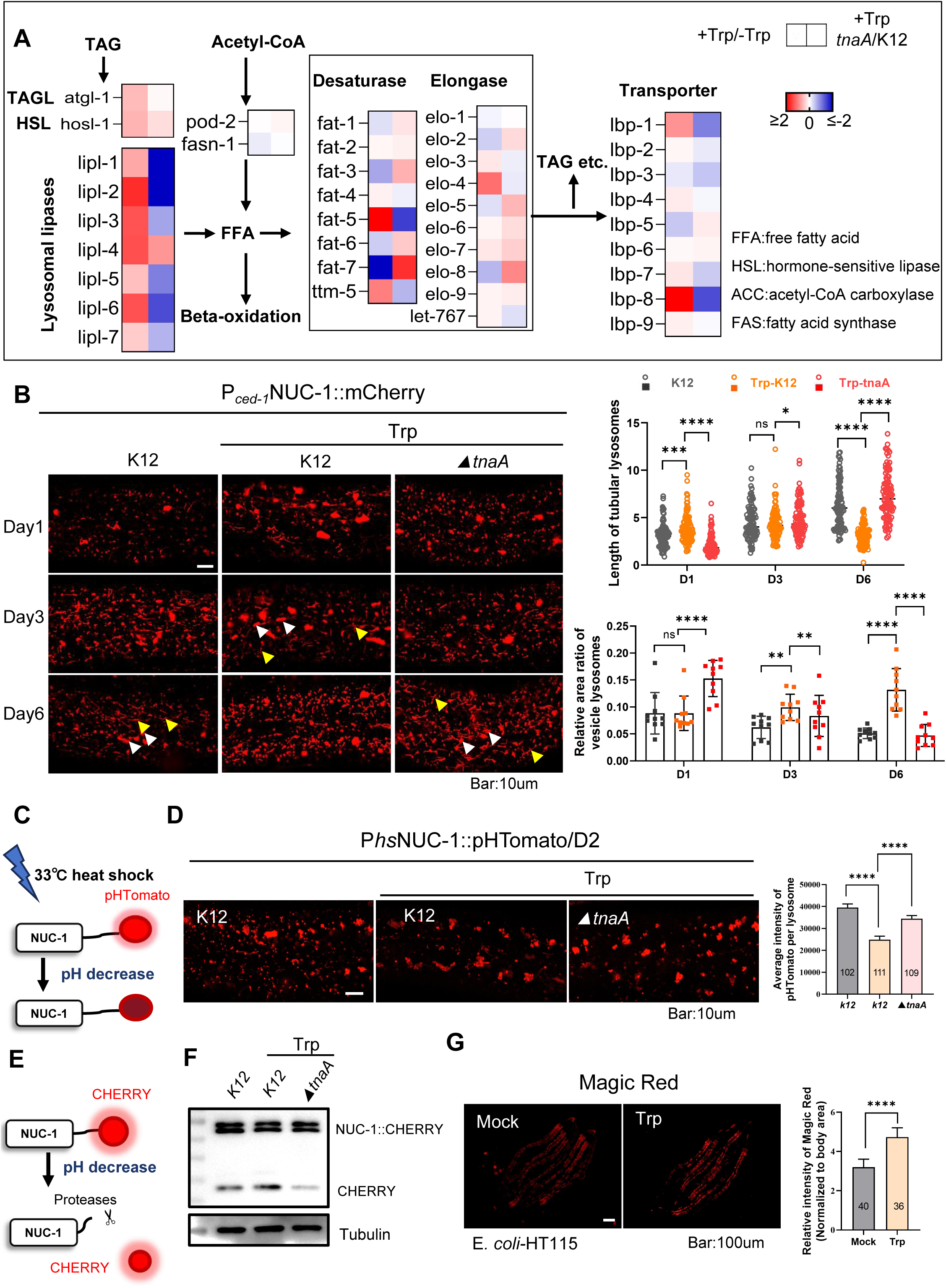
Bacterial tryptophan metabolism activates lysosomal function. **(A)** Heat map showing fold changes in mRNA levels of lipid-metabolism genes in worms fed Trp-supplemented K12 versus mock-treated K12, and Trp-supplemented Δ*tnaA* versus Trp-supplemented K-12. Fold changes were calculated by dividing each gene’s expression level under the indicated conditions. Source data are provided in Table S1. **(B)** Confocal images of lysosomal morphology in wild-type adults expressing NUC-1::mCherry at days 1, 3, and 6 of adulthood. Animals were maintained on NGM plates seeded with K12 (control) or with K12/Δ*tnaA* plus 10 mM tryptophan. White arrowheads denote vesicular lysosomes; yellow arrowheads indicate distinct lysosomal tubules. The relative area of vesicular lysosomes and the length of per lysosomal tubules were quantified per unit area 31×43 um^2^. (the relative rare of vesicular lysosome=Sum of vesicular lysosome areas in unit area/unit area 31×43 um^2)^) .At least 10 worms were scored in each group at each age. Scale bar, 10 µm. **(C)** Schematic of the heat-shock-inducible NUC-1::pHTomato reporter. Fusion of the lysosomal nuclease NUC-1 to the pH-sensitive fluorophore pHTomato allows real-time monitoring of lysosomal pH. Fluorescence decreases as lysosomal acidification increases (i.e., as pH declines). **(D)** Confocal micrographs and quantitative analysis of lysosomes in day-2 adults expressing hsNUC-1::pHTomato after heat shock. Animals were maintained on NGM plates seeded with K12 (control) or with K12/Δ*tnaA* plus 10 mM tryptophan. The average intensity of pHTomato per lysosome is shown in the right. At least 20 animals were scored in each group. Scale bar, 10 µm. **(E)** Diagram of the lysosomal degradation assay based on cleavage of NUC-1::mCherry in worms. **(F)** Western blot of mCherry cleavage products from NUC-1::mCherry in day-1 adults fed K12 or K-12/Δ*tnaA* with tryptophan, demonstrating enhanced lysosomal degradation. **(G)** Representative fluorescence images (left) and quantification (right) of Magic Red cathepsin activity in L4 worms fed on NGM plate with or without 10 mM tryptophan, seeded with E. coli-HT115. The number of animals analyzed is indicated. Scale bar, 100 µm. Data represent mean ± SD. All statistical analyses were performed using unpaired two-tailed Student’s t-test. ****p < 0.0001. All experiments were performed independently at least three times. See also Figure S3 and Table S1.

KEGG pathway analysis of differentially expressed genes (Trp-K12 vs Mock-K12) underscored the lysosome as a key pathway responsive to bacterial tryptophan metabolites (Figure S3C), with 14 lysosome-related genes (Figure S3D)—including various lipohydrolases and other hydrolases—being significantly induced at both group (Trp-K12 vs Mock-K12, Trp-K12 vs Trp-tnaA)(Figure S3E, S3F). Given that lysosomes play a critical role in maintaining lipid metabolism by breaking down and recycling various lipid species. They contain a variety of acid hydrolases—including lysosomal acid lipase—that digest lipids such as triglycerides, cholesterol esters, and other complex lipids into free fatty acids and cholesterol (Settembre and Ballabio, 2014a). Thus, it is possible that bacterial tryptophan metabolism regulates lysosomal function to mediates lipid metabolism in *C. elegans*.

### Bacterial Tryptophan Metabolism Activates Lysosomal Function

To investigate whether bacterial tryptophan metabolism modulates lysosomal activity, we systematically evaluated lysosomal morphology, acidification, and degradation capacity in *C. elegans* fed *E. coli* K12 or tnaA mutants under tryptophan-supplemented conditions.

First, we tracked age-associated lysosomal morphology changes using the established reporter (XW5399:*qxIs257*[P*_ced-1_*NUC-1::CHERRY]) (Miao et al., 2020; Sun et al., 2020) fed K12 (Mock-K12), K12 with tryptophan (Trp-K12), or tnaA mutants with tryptophan (Trp-tnaA). Consistent with prior reports (Sun et al., 2020), Mock-K12 animals (grown on K12-*E. coli* without tryptophan) exhibited age-dependent lysosomal remodeling, including reduced vesicular structures and increased tubular morphology. In contrast, Trp-K12 animals (fed K12-*E. coli* with tryptophan) maintained a shorter tubular lysosomes and hihger proportion of vesicular lysosomes through aging (at Day 3 and Day 6), but not at Day 1 (Figure 3B). Strikingly,

Trp-tnaA animals (fed tnaA mutant *E. coli* with tryptophan) mirrored Mock-K12 morphology, by increasing tubular lysosomes and reduceing vesicular structures with aging (Figure 3B). This data indicates that bacterial tryptophan metabolism preserves a youthful lysosomal architecture .

Next, we assessed lysosomal acidification using the transgenic reporter NUC-1::pHTomato (XW19180: *qxIs750*[P*_hs_*NUC-1::pHTomato]) (Sun et al., 2020). In this system, the pH-sensitive fluorescent protein pHTomato (pKa ≈7.8) (Li and Tsien, 2012) is fused to NUC-1 and expressed under a heat-shock promoter; pHTomato decreases in fluorescence at lower pH values (Figure 3C). We found that Trp-K12 animals (fed K12-*E. coli* with tryptophan) exhibited significantly lower NUC-1::pHTomato fluorescence compared to Mock-K12 animals (grown on K12-*E. coli* without tryptophan) (Figure 3D). Importantly, this reduction in fluorescence was reversed in Trp-tnaA animals (fed tnaA mutant *E. coli* with tryptophan) (Figure 3D), indicating that bacterial tryptophan metabolism enhances lysosomal acidification—a condition that is critical for the optimal activity of lysosomal hydrolases.

Finally, we evaluated lysosomal degradation activity using the transgenic reporter-*qxIs257,* P*_ced-1_*::NUC-1::CHERRY (Miao et al., 2020). When NUC-1::CHERRY delivered to lysosomes, CHERRY is cleaved by cathepsins, and the cleaved products can be quantified by western blot (Figure 3E). Our results showed that, following tryptophan supplementation in *E. coli* K12, there was a induction in cleaved CHERRY levels; in contrast, no such induction was observed in animals fed the tnaA mutant (Figure 3F). Additionally, using a cathepsin assay with MagicRed (Pu and Qi, 2024)—a substrate that emits red fluorescence upon cleavage by Cathepsin B—we observed increased lysosomal degradation activity in animals fed *E. coli* HT115 with tryptophan supplementation (Figure 3G). This data indicates that bacterial tryptophan metabolism increases lysosomal degradation activity in *C. elegans*.

In summary, our data demonstrate that bacterial tryptophan metabolism enhances lysosomal function in *C. elegans* by (1) promoting a more youthful lysosomal morphology, (2) increasing lysosomal acidification, and (3) boosting lysosomal degradation activity.

### Bacterial Tryptophan Metabolism Drives Lysosomal-Dependent Lipid Degradation

To test whether bacterial tryptophan metabolism reduces host lipid content by enhancing lysosomal activity, we genetically disrupted lysosomal function and assessed its impact on *lbp-8* expression and lipid levels.

First, we targeted *cup-5*, encoding a lysosomal Ca²⁺ channel homologous to human TRPML that regulates lysosomal acidity and activity (Hersh et al., 2002; Miao et al., 2020; Sun et al., 2011). Knockdown of *cup-5* in Trp-K12 animals (fed tryptophan-supplemented *E. coli* K12) abolished the tryptophan-induced reduction in lipid content (Figure 4A) and suppressed *lbp-8* upregulation (Figure S4A–S4B). Similarly, knockdown of *lipl-4*—a lysosomal acid lipase required for lipid hydrolysis at low pH (Folick et al., 2015)—restored lipid levels (Figure 4A) and attenuated *lbp-8* induction in Trp-K12 animals (Figures S4A-S4B). These findings demonstrate that bacterial tryptophan metabolism drives lipid degradation through lysosomal acidification and lipase activation.

**Figure 4.**
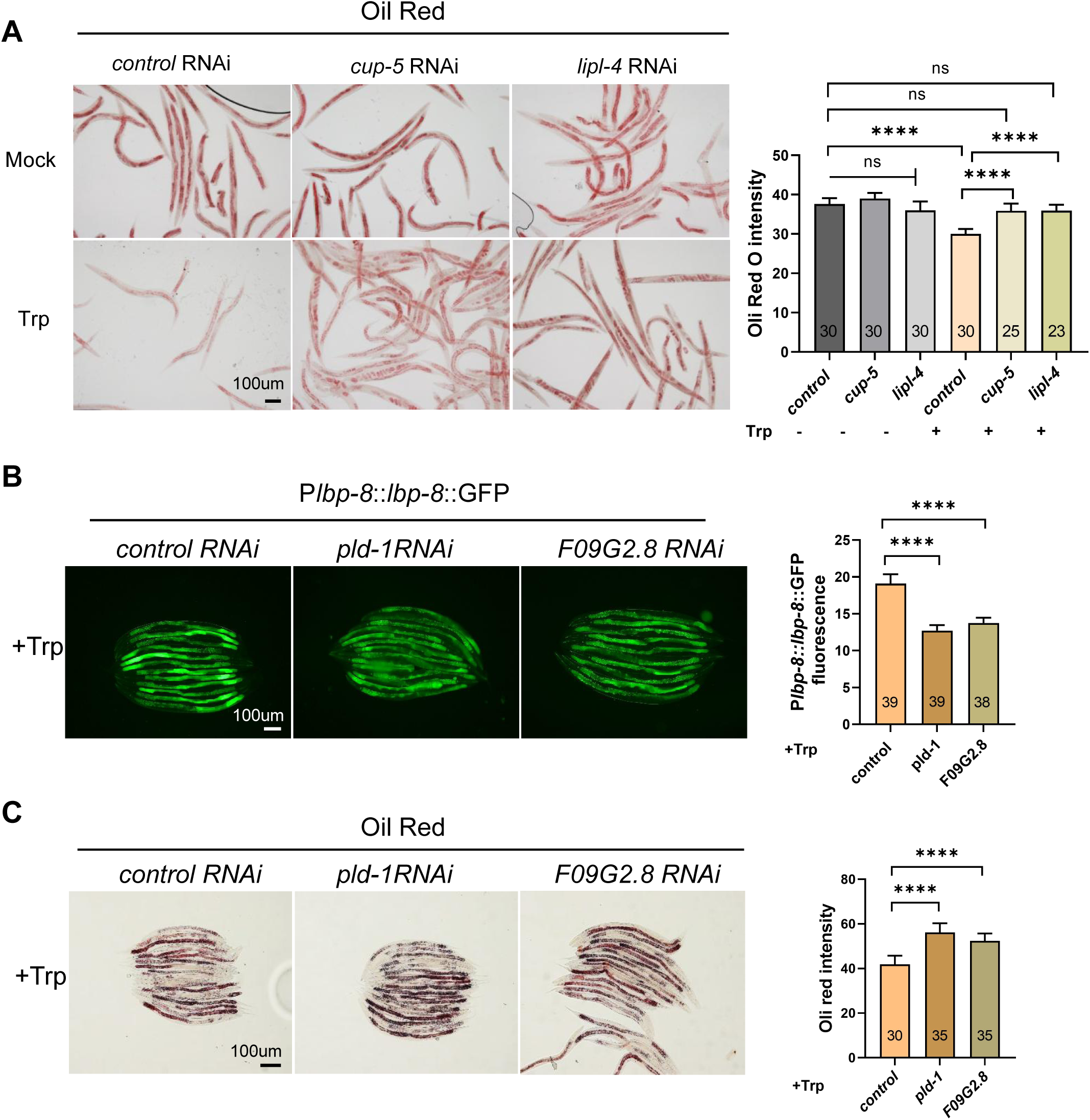
Bacterial tryptophan metabolism promotes lysosomal-dependent lipid degradation. **(A)** Oil Red O staining and quantification of total lipid levels in wild-type L4 worms treated with control, *cup-5*, or *lipl-4* RNAi, grown on NGM plates with or without 10 mM tryptophan. Scale bar, 100 µm. **(B)** Representative fluorescence images and quantification of the P*lbp-8*::GFP reporter in wild-type L4 worms with control, *pld-1*, or *F09G2.8* RNAi, cultured on NGM + 10 mM tryptophan. Scale bar, 100 µm. **(C)** Oil Red O staining and quantification of lipid content in wild-type L4 worms with control, *pld-1*, or *F09G2.8* RNAi, cultured on NGM + 10 mM tryptophan. Scale bar, 100 µm. Data represent mean ± SD. The number of animals analyzed is indicated. ****p < 0.0001 (ANOVA with multiple-comparisons correction). All experiments were performed independently at least three times. See also Figure S4.

Lysosomal lipid degradation—whether of lipoproteins, lipid droplets, or cell membranes—occurs predominantly within the lysosomal lumen and depends on the formation of intraluminal vesicles (ILVs). A specific phospholipid, bis(monoacylglycero)phosphate (BMP), is critical for ILV formation and BMP-mediated lipid degradation (Frederick et al., 2009; Singh et al., 2024). Previous studies have reported that lysosomal enzymes PLD3 and PLD4 are necessary for maintaining normal BMP levels in human cells and murine tissues (Singh et al., 2024), thereby regulating lipid degradation in lysosomes. We found that knockdown of F09G2.8 (homologous to PLD3/PLD4) or *pld-1* (a phospholipase D homolog) (Figure S4C) partially reversed the activation of *lbp-8* and the reduction in lipid content in Trp-K12 animals (Figures 4B-4C). This demonstrates that bacterial tryptophan metabolism relies on BMP-dependent ILV formation to promote lysosomal lipid degradation.

Collectively, these findings indicate that bacterial tryptophan metabolism promotes lipid degradation in *C. elegans* through functional lysosomes.

### Bacterial Tryptophan Metabolism Enhances Mitochondrial β-oxidation through Lysosomal Activation to Promote Lipid Metabolism

To identify regulatory factors driving increased lipid mobilization in *C. elegans* exposed to bacterial tryptophan metabolites, we conducted gene ontology (GO) enrichment analysis of 1334 genes being significantly induced at both condition (Trp-K12 vs Mock-K12, Trp-K12 vs Trp-tnaA) (Figure S3B). Go enrichment showed significant enrichment in categories such as “monooxygenase activity”, and “oxidoreductase activity” which involved in mitochondria metabolsim (Figure S5A).

During lipid catabolism, lysosomal lipases hydrolyze stored triglycerides into free fatty acids (FFAs), which are subsequently processed via mitochondrial and peroxisomal β-oxidation to generate ATP (Grabner et al., 2021). Key genes involved in fatty acid β-oxidation—including acyl-CoA synthetases (ACS) and acyl-CoA dehydrogenases (ACDH)—were upregulated in Trp-K12-fed animals compared to controls (Figure S5B).

To directly assess β-oxidation activation, we generated transgenic worms expressing *acs-2*::GFP under the control of its native promoter (P*acs-2*). We found that expression of P*acs-2:: acs-2*::GFP is increased in Trp-K12-fed animals but not in those fed Trp-tnaA mutants (Figure 5A), indicating bacterial tryptophan metabolism specifically induces *acs-2*.

**Figure 5.**
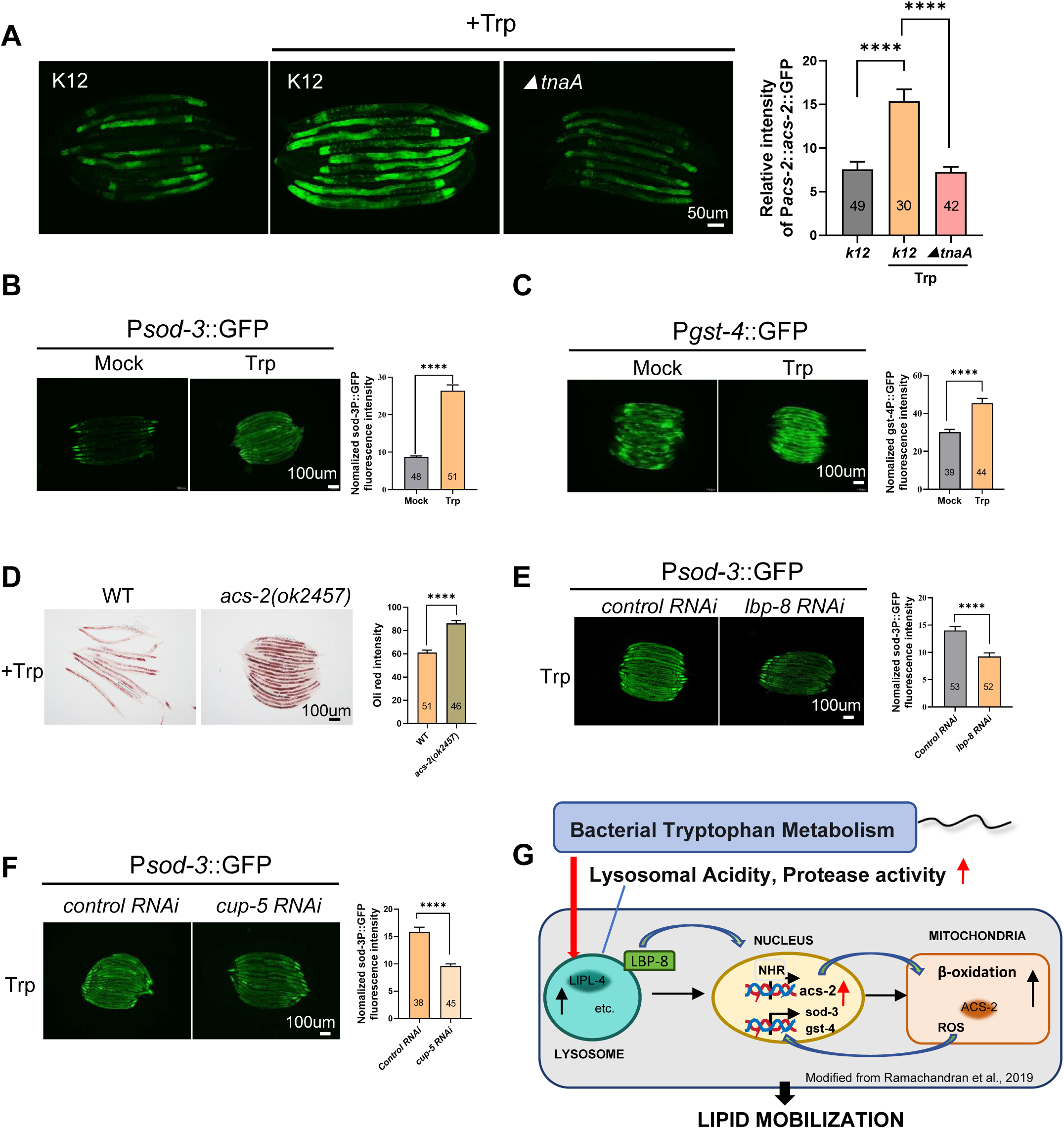
Bacterial tryptophan metabolism enhances mitochondrial β-oxidation via lysosomal activation. **(A)** Representative fluorescence images and quantification (right) of the P*acs-2::acs-2::*GFP reporter in L4 worms fed on NGM plates seeded with wild-type K12 (control) or with K-12/Δ*tnaA* plus 10 mM tryptophan for 60 h. The number of animals analyzed is indicated. Scale bar, 50 µm. **(B)** Representative images and quantification of P*sod-3::*GFP in L4 worms grown on NGM with or without 10 mM tryptophan, seeded with *E. coli*-K12. The number of animals analyzed is indicated. Scale bar, 100 µm. **(C)** Representative images and quantification of P*gst-4::*GFP in L4 worms grown on NGM with or without 10 mM tryptophan, seeded with *E. coli*-K12. The number of animals analyzed is indicated. Scale bar, 100 µm. **(D)** Oil Red O staining and quantification of lipid content in wild-type and *acs-2(ok2457)* mutant L4 worms fed NGM + 10 mM tryptophan, seeded with *E. coli*-K12. The number of animals analyzed is indicated. Scale bar, 100 µm. **(E)** Representative images and quantification of P*sod-3::*GFP in worms subjected to control or *lbp-8* RNAi on NGM + 10 mM tryptophan. Scale bar, 100 µm. **(F)** Representative images and quantification of P*sod-3::*GFP in worms subjected to control or *cup-5* RNAi on NGM + 10 mM tryptophan. The number of animals analyzed is indicated. Scale bar, 100 µm. **(G)** Model of the lysosome–LBP-8–mitochondrial signaling axis by which bacterial tryptophan metabolism stimulates lipid β-oxidation through lysosomal activation. Previous studies have shown that lysosomal signaling extends lifespan by modulating mitochondrial activity—particularly through the LIPL-4–LBP-8 pathway, which upregulates mitochondrial β-oxidation to promote lipid metabolism (Ramachandran et al., 2019) Data represent mean ± SD. All statistical analyses were performed using unpaired two-tailed Student’s t-test. ****p < 0.0001. All experiments were performed independently at least three times. See also Figure S5.

Enhanced mitochondrial or peroxisomal β-oxidation often elevates reactive oxygen species (ROS), byproducts of oxidative phosphorylation. Using the redox-sensitive dye MitoTracker™ Red CMXRos—which reflects mitochondrial membrane potential and ROS levels (Li et al., 2019; Liu et al., 2024)—we observed significantly increased ROS in Trp-K12-fed animals (Figure S5C). This effect was absent in tnaA mutant-fed animals (Figure S5C). Consistent with this, oxidative stress-responsive genes, including *sod-3* (superoxide dismutase; Figure 5B) and *gst-4* (glutathione S-transferase; Figure 5C), were upregulated in Trp-K12-fed worms. Crucially, *acs-2* knockout abolished the enhanced lipid degradation in Trp-K12-fed animals (Figure 5D), demonstrating that ACS-2-dependent mitochondrial β-oxidation is required for tryptophan-mediated lipid loss.

Notably, RNAi knockdown of lysosomal genes *cup-5* or *lbp-8* under tryptophan supplementation reduced mitochondrial ROS levels, as measured by MitoTracker™ Red CMXRos (Figures S5E, S5F) and the oxidative stress reporter P*sod-3*::GFP (Figures 5E, 5F).

Previous studies have shown that lysosomal signaling extends lifespan by modulating mitochondrial activity — particularly through the LIPL-4 – LBP-8 pathway, which upregulates mitochondrial β-oxidation to promote lipid metabolism (Ramachandran et al., 2019; Figure 5G). Our findings now uncover a novel mechanism: bacterial tryptophan metabolism stimulates lysosomal activity, thereby enhancing mitochondrial β-oxidation. This cascade not only boosts lipid catabolism but also increases reactive oxygen species (ROS) production.

### Bacterial Tryptophan Metabolism Enhances Lysosomal Function Which Promotes Lipid Breakdown in Liver Cells

To investigate the conservation of the mechanism by which bacterial tryptophan metabolism activates lysosomal function to promote lipolysis, we first established an experimental model using the human liver cell line Huh7. Cells were treated with either *E. coli* culture supernatants grown in LB medium supplemented with tryptophan (Trp-K12-sup), supernatants from LB medium without tryptophan (K12-sup), or LB medium alone (control) (Figure 6A).

**Figure 6.**
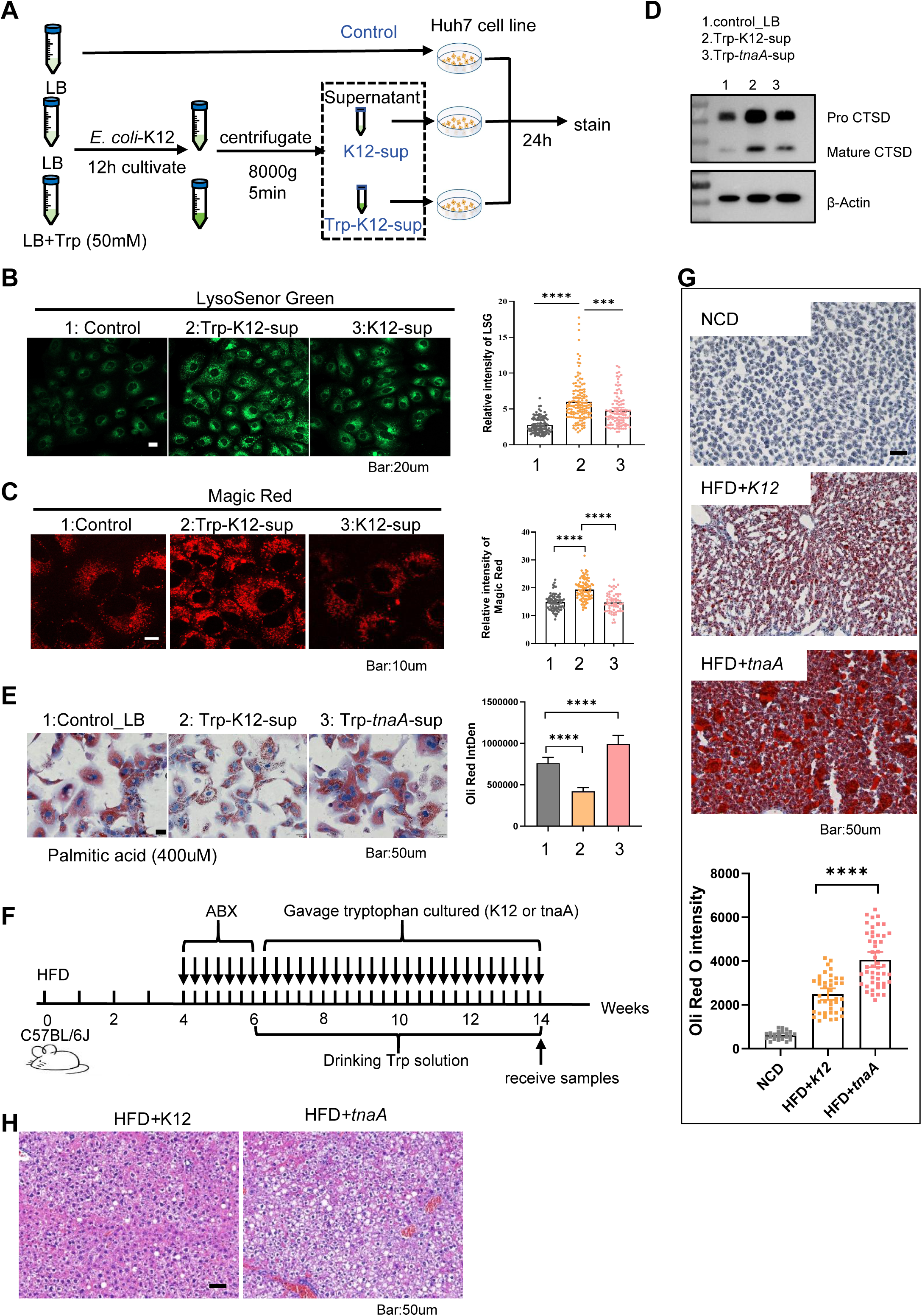
Bacterial tryptophan metabolism activates lysosomal function in mammalian cells and promotes lipid breakdown in a high-fat diet mouse model. **(A)** Schematic of in vitro treatments: Huh7 cells were incubated with *E. coli* K12 culture supernatants prepared with tryptophan supplementation (Trp-K12-sup), without tryptophan (K12-sup), or LB medium alone (LB). **(B)** Confocal micrographs and quantification of LysoSensor™ Green DND-189 fluorescence in Huh7 cells treated as in (A). Scale bar, 20 µm. **(C)** Confocal micrographs and quantification of Magic Red cathepsin activity in Huh7 cells treated as in (A).Scale bar, 10 µm. **(D)** Western blot analysis of endogenous cathepsin D (CTSD) in Huh7 cells treated with tryptophan-supplemented K-12 supernatant (Trp_k12 sup), tryptophan-supplemented Δ*tnaA* supernatant (Trp_tnaA sup), or LB control. Both precursor and mature CTSD bands are shown. **(E)** Representative Oil Red O staining and quantification of lipid accumulation in primary B6J hepatocytes co-treated for 24 h with 400 µM palmitic acid and supernatants from Trp-supplemented K-12/Δ*tnaA, or* LB medium alone (LB).Scale bar, 50 µm. **(F)** Experimental timeline for the high-fat diet (HFD) mouse study: mice were fed HFD and received oral gavage of Trp-supplemented K12 or Δ*tnaA* cultures as indicated. See more detail at methods: Animal experiments and procedures. **(G)** Representative Oil Red O–stained liver sections from HFD-fed mice as shown in (F) and quantification of staining intensity. Scale bar, 50 µm. **(H)** Representative hematoxylin & eosin–stained liver sections from HFD-fed mice as shown in (F) . Scale bar, 50 µm. Data represent mean ± SD. All statistical analyses were performed using unpaired two-tailed Student’s t-test. ****p < 0.0001, ***p < 0.001. All experiments were performed independently at least three times. See also Figure S6.

To investigate the acidification of lysosomes, Hub7 cells were stained with LysoSensor^TM^ Green DND-189 as the dye accumulates in lysosomes resulting in an increased fluorescence intensity upon acidification. The supernatant from Trp-K12-sup (tryptophan-supplemented *E. coli*) cultures significantly increased lysosomal acidity compared to K12-sup (supernatant from *E. coli*) or control (only LB medium). (Figue 6B) This indicates that bacterial tryptophan metabolism increased lysosomal acidity in liver cells.

Next, we evaluated lysosomal degradation activity using MagicRed staining—a substrate that emits red fluorescence upon cleavage by Cathepsin B. The supernatant from Trp-K12-sup (tryptophan-supplemented *E. coli*) cultures significantly increased red fluorescence compared to K12-sup (supernatant from *E. coli*) or control (only LB medium), suggesting bacterial tryptophan metabolism increased lysosomal degradation activity. Cysteine cathepsins are responsible for driving proteolytic degradation within the lysosome (Man and Kanneganti, 2016; McGlinchey and Lee, 2015). Western blot analysis of endogenous cathepsin D (CTSD) processing revealed elevated levels of both precursor and mature forms in cells treated with tryptophan-containing supernatants from *E. coli*-K12 (Trp-K12-sup), but not from *E. coli*-tnaA mutant (Trp-tnaA-sup) (Figure 6D). This data further indicated that bacterial-tnaA mediated tryptophan metabolism increased lysosomal degradation activity in liver cells.

To evaluate whether bacterial tryptophan metabolism promote lipid breakdown, primary mouse hepatocytes were co-treated with palmitic acid and supernatants from *E. coli* K12 (Trp-K12-sup) or tnaA mutant (Trp-tnaA-sup) cultures. Oil Red O staining revealed significantly reduced lipid accumulation in cells exposed to K12 supernatants, whereas tnaA mutant supernatants showed no difference from controls (Figure 6E), indicating the role of bacterial tryptophan metabolism in enhancing lipid metabolism in liver cells.

Collectively, our data demonstrate that bacterial tryptophan metabolism enhances lysosomal activity, promoting lipid degradation in hepatocytes.

### Bacterial Tryptophan Metabolism Promotes Lipid Breakdown in High-Fat Diet Mice

We further explored the correlation between bacterial tryptophan metabolism and host lipid content in vivo using an obesity mouse model which was established by feeding mice a high-fat diet (HFD) for 14 weeks (Figure 6F). The mice were then treated with antibiotics for 2 weeks (from week 4 to week 6) to eliminate intestinal bacteria. The universal bacterial 16S rRNA qPCR data indicated that the majority of gut microbiota were eliminated after antibiotics treatment by using faeces as sample (Figure S6A).

From week 6 to week 14, animals were subjected to oral gavage with either wild-type *E. coli* K12 or tnaA mutant strains cultured with tryptophan, co-administered with tryptophan supplementation (Figure 6F).

Although the body weights of the two groups did not differ significantly over time (Figure S6B), the liver weight of mice treated with bacterial tryptophan metabolites was reduced (Figure S6D). Oil red O staining of liver sections revealed that the HFD group receiving the tnaA mutant (lacking effective tryptophan metabolism) exhibited more red-stained lipid droplets compared to the group receiving the wild-type K12 strain (Figure 6G). Hematoxylin and eosin (HE) staining further confirmed that the HFD-tnaA group contained numerous fat vacuoles relative to the HFD-K12 group (Figure 6H).

In summary, our data demonstrate that bacterial tryptophan metabolism activates lysosomal function in mammalian cells, leading to increased lipid degradation. In a high-fat diet mouse model, enhancing intestinal bacterial tryptophan metabolism alleviates liver fat accumulation.

## Discussion

In this study, we reveal a conserved mechanism whereby gut bacterial tryptophan metabolism activates host lysosomal function to facilitate lipid breakdown (Figure 7). We demonstrated that *E. coli* TnaA-mediated tryptophan metabolism enhances lysosomal acidification, enzymatic degradation capacity, and structural integrity, leading to induction of the lipid chaperone LBP-8, increased mitochondrial β-oxidation, and reduced lipid storage. Importantly, these findings extend to mammalian hepatocytes and a high-fat diet mouse model, where restoration of bacterial tryptophan metabolism alleviates hepatic steatosis. Our work uncovers microbiota-regulated lysosomal activation as a critical axis in lipid homeostasis and highlights its potential as a therapeutic target for metabolic disorders linked to lysosomal dysfunction.

**Figure 7.**
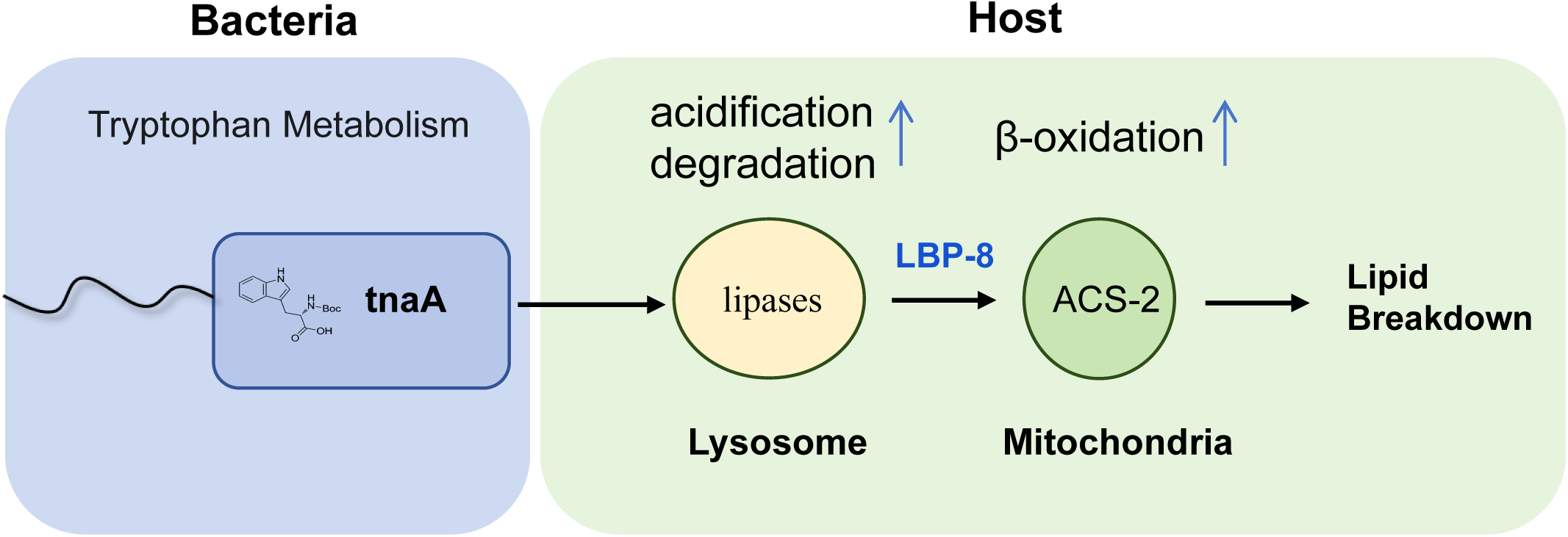
A proposed model shows that bacterial *tnaA*-mediated tryptophan metabolism enhances lysosomal function, leading to LBP-8 expression and activation of β-oxidation to facilitate lipid degradation.

The interaction between gut microbiota and lysosomes differs from the evasion strategies employed by intracellular pathogens. Intracellular pathogens such as *Mycobacterium tuberculosis* and *Salmonella* subvert host lysosomal pathways to evade immune defenses by impairing acidification and trafficking (Sreelatha et al., 2015; Wong et al., 2011; Xu et al., 2019). In contrast, the ability of commensal gut bacteria to modulate lysosomal function has remained unexplored. Our identification of bacterial tryptophan metabolism as a potent activator of host lysosomes fills a critical gap in our understanding of how the gut microbiome influences fundamental cellular processes. This suggests a symbiotic relationship where gut microbes optimize lysosomal function to maintain metabolic homeostasis, contrasting with pathogenic subversion of lysosomal pathways. Such duality underscores the importance of understanding how microbial metabolites differentially regulate lysosomal dynamics in health and disease.

Clinical and preclinical studies reveal that metabolic syndrome is associated with diminished microbial production of AhR agonists, leading to reduced activation of AhR-dependent pathways that are essential for metabolic health (Natividad et al., 2018). This deficiency in AhR ligand production directly correlates with disease severity, highlighting a causal link between disrupted bacterial tryptophan metabolism and metabolic dysfunction. Despite these advances, how microbial tryptophan metabolites interface with lysosomal pathways to regulate host metabolism (lipid homeostasis) remained unexplored. Our discovery that bacterial tryptophan metabolism activates lysosomal function, a key regulator of lipid catabolism, and that this activation promotes lipid breakdown in both cellular and animal models provides a mechanistic link between microbial tryptophan metabolism and host metabolic health. This suggests that restoring or enhancing bacterial tryptophan metabolism could be a promising therapeutic strategy to reactivate lysosomal lipolysis and ameliorate lipid-related pathologies.

Beyond lipid metabolism, lysosomes are increasingly recognized for their diverse roles in cellular processes, including nutrient sensing, autophagy, aging, and tissue regeneration (Folick et al., 2015; Miao et al., 2020; Settembre and Ballabio, 2014a). Given the broad involvement of lysosomes in maintaining cellular homeostasis, the finding that bacterial tryptophan metabolism can activate these organelles suggests potential implications beyond lipid metabolism. Future studies could explore whether this pathway plays a role in other physiological contexts regulated by lysosomal function, such as longevity, inflammation, and neurodegenerative diseases.

Investigating the specific tryptophan metabolites responsible for lysosomal activation and the precise host cell receptors or signaling pathways involved will further illuminate this crucial host-microbe interaction and potentially uncover novel therapeutic targets for a wide range of human diseases.

### Limitations of the Study

While our study establishes that bacterial tryptophan metabolism activates lysosomal function to promote lipid breakdown, several limitations merit attention. First, the molecular mechanisms by which host organisms’ sense bacterial tryptophan metabolites to activate lysosomal pathways remain unresolved. We propose future genome-wide genetic screens using the P*lbp-8*::GFP reporter to identify host factors mediating this recognition. Second, whether specific tryptophan-derived metabolites (e.g., indole) directly target lysosomes to modulate their activity remains unverified; metabolomic profiling of lysosomal compartments could clarify this interaction. Third, while lysosomal activation was inferred indirectly via lipid clearance in mice, technical limitations precluded direct measurement of lysosomal acidification in hepatic tissues. Development of pH reporters (e.g., NUC-1::pHTomato) for in vivo lysosomal tracking in murine models will bridge this gap. Addressing these gaps will elucidate the molecular mechanisms through which host organisms detect bacterial tryptophan metabolites and orchestrate lysosomal functional activation, thereby advancing our understanding of microbiota-regulated metabolic homeostasis.

## Resource availability

### Materials availability

All reagents and strains generated by this study are available through request to the corresponding author (Zhao Shan; Bin Qi) with a completed Material Transfer Agreement.

### Data and code availability

- RNA-seq data are provided in Table S1
- This paper does not report original code.
- Any additional information required to reanalyze the data reported in this paper is available from the corresponding author (Zhao Shan; Bin Qi) upon request.

## Acknowledgments

We thank the Caenorhabditis Genetics Center (CGC) (funded by NIH P40OD010440) for strains; Dr. Xiaochen Wang (SUSTech) and Chonglin Yang (Yunnan U.) for sharing strains and suggestions. This work was supported by the Yunnan Provincial Science and Technology Project at Southwest United Graduate School (202302AP370005 to B.Q.), National Natural Science Foundation of China (32170794 to B.Q.; 32071129 to Z.S), Yunnan Revitalization Talent Support Program (C619300A086 to Z.S., K264202230211 to B.Q.).

## Author Contributions

K.Z and Z.L performed most experiments. Y.L performed the *E. coli* keio screen. Y.C, L.W and Y.L performed some mice experiment with K.Z. J.Z performed some RNAi screen. Q.L and R.Y construct some transgenic animals. Z.S, B.Q and K.Z. wrote and edited the paper. Z.S and B.Q supervised this study.

## Supplemental Figure Legends

**Figure S1. P*lbp-8::*GFP reporter expression on metabolically compromised *E. coli, and* Bacterial growth state on NGM and LB plate. Related to Figure 1**.

**(A)** Representative fluorescence images of the *Plbp-8::*GFP reporter in worms fed with metabolically compromised *E. coli* (treated with ampicillin or UV-killed). “ns” indicates no significant difference (p > 0.05, Student’s *t*-test). Data represent mean ± SD. The number of analyzed animals is indicated. Scale bar: 100 µm.

**(B)** Bacterial growth state on NGM and LB culture media seed with *E. coli*-K12.

**Figure S2. Bacterial tryptophan metabolism induce *lbp-8* expression. Related to Figure 2**.

**(A)** Representative fluorescence images and quantification of the P*lbp-8::*GFP reporter in worms fed with *E. coli* K12 mutants, which significantly suppressed *lbp-8* expression on LB medium. Scale bar: 100µm.

**(B)** *lbp-8* mRNA levels in L4-stage wild-type worms exposed to *E. coli* K12 on NGM (NGM-K12) and *E. coli* K12 or *tnaA* cultured on LB medium (LB-K12/LB-*tnaA*), which was extracted from RNA-seq data. Data represent mean ± SD. ****p < 0.0001, Student’s *t*-test.

**(C)** Fluorescence images of the P*lbp-8::lbp-8::*GFP reporter in worms exposed to wild-type *E. coli* K12 or *tnaA* mutants on LB medium, along with quantification of nucleus-to-cytoplasm GFP intensity of LBP-8 protein. Data represent mean ± SD. ****p < 0.0001, Student’s *t*-test. The number of animals analyzed is indicated. Scale bar: 10 µm.

**(D)** *lbp-8* mRNA levels in L4-stage wild-type worms exposed to *E. coli* K12 grown under standard NGM conditions (Mock) or supplemented with 10 mM tryptophan (*E. coli* K12-Trp, *tnaA*-Trp), which was extracted from RNA-seq data. Data represent mean ± SD. ****p < 0.0001, Student’s *t*-test.

**(E)** Fluorescence images of the P*lbp-8::lbp-8::*GFP reporter in worms exposed to wild-type *E. coli* on NGM medium with or without 10 mM tryptophan, along with quantification of nucleus-to-cytoplasm GFP intensity of LBP-8 protein. Data represent mean ± SD. ****p < 0.0001, Student’s *t*-test. The number of animals analyzed is indicated. Scale bar: 10 µm.

**(F)** Fluorescence images of the P*lbp-8::lbp-8::*GFP reporter in worms exposed to wild-type *E. coli* on NGM medium with or without 10 mM tryptophan. Data represent mean ± SD. ****p < 0.0001, Student’s *t*-test. The number of animals analyzed is indicated. Scale bar: 100 µm.

**(G)** Fluorescence images of the P*lbp-8::lbp-8::*GFP reporter in worms exposed to wild-type *E. coli* or heat-killed *E. coli* cultured in LB medium with or without tryptophan. Data represent mean ± SD. ****p < 0.0001, ns indicates no significant difference (p > 0.05, Student’s *t*-test). The number of animals analyzed is indicated. Scale bar: 100 µm.

**Figure S3. Bacterial tryptophan metabolism induce lysosomes-related genes expression. Related to Figure 3**.

**(A)** Schematic diagram illustrating the RNA sequencing sample preparation process, including the three experimental conditions: NGM plates seeded with *E. coli* K12 (Mock-K12), tryptophan-supplemented NGM plates seeded with *E. coli* K12 (Trp-K12), and tryptophan-supplemented NGM plates seeded with *E. coli tnaA* (Trp-*tnaA*). Synchronized L1-stage wild-type animals were grown on these plates until reaching the L4 stage, at which point L4-stage animals were collected for sequencing.

**(B)** Venn diagram showing the overlap between genes differentially expressed in Trp-K12 vs NGM-K12 and in Trp-K12 vs Trp-*tnaA*.

**(C)** KEGG enrichment analysis of differentially expressed genes in Trp-K12 vs NGM-K12.

**(D)** Venn diagram displaying the overlap of lysosomal-related genes, which are induced at both group (Trp-K12 vs Mock-K12, Trp-K12 vs Trp-tnaA).

**(E)** List of overlapping lysosomal-related genes.

**(F)** mRNA levels from RNA-seq of the lysosomal-related genes identified in (D).

**Figure S4. Bacterial tryptophan metabolism promotes lipid degradation in a lysosomal-dependent manner. Related to Figure 4**.

**(A-B)** Representative fluorescence images (A) and quantification (B) of the P*lbp-8*::GFP reporter in wild-type L4 worms treated with control, *cup-5*, *lipl-4* or *lbp-8* (negative control) RNAi, grown on NGM plates with or without 10 mM tryptophan. Data represent mean ± SD. The number of animals analyzed is indicated. ****p < 0.0001, “ns” p > 0.05 (Student’s t-test). Scale bar, 100 µm.

**(C)** Homology alignment of *C. elegans* PLD-1 and F09G2.8 with human PLD-3 and PLD-4 protein sequences. For each row/column, 100% represents the target protein, and percentages indicate sequence similarity relative to the target protein.

**Figure S5. Bacterial tryptophan metabolism enhances mitochondrial β-oxidation via lysosomal activation to facilitate lipid metabolism. Related to Figure 5**.

**(A)** Gene ontology (GO) enrichment analysis of 1334 genes being significantly induced at both condition (Trp-K12 vs Mock-K12, Trp-K12 vs Trp-tnaA) (Figure S3B).

**(B)** Heatmap showing fold changes in mRNA levels of β-oxidation-related genes in worms fed Trp-supplemented K12 (Trp-K12) versus mock-treated K12 (Mock-K12), and Trp-supplemented Δ*tnaA* (Trp-tnaA) versus Trp-supplemented K-12 (Trp-K12). Fold changes were calculated by dividing each gene’s expression level under the indicated conditions.

**(C)** Representative fluorescence images and quantification of MitoTracker™ Red CMXRos in L4 worms fed on NGM plates seeded with wild-type K12 (control) or with K12/Δ*tnaA* plus 10 mM tryptophan. Data represent mean ± SD. The number of animals analyzed is indicated. **p < 0.01, *p < 0.05 (Student’s t-test). Scale bar: 100 μm.

**(D-E)** Representative microscope images and quantification of MitoTracker™ Red CMXRos fluorescence in wild-type, *acs-2(ok2457)* (D) or *lbp-8(gk5151)* (E) mutants L4 worms fed NGM + 10 mM tryptophan, seeded with *E. coli*-K12. Data represent mean ± SD. The number of animals analyzed is indicated. ****p < 0.0001 (Student’s t-test). Scale bar: 100 μm.

**(F)** Representative microscope images and quantification of MitoTracker™ Red CMXRos fluorescence in wild-type worms subjected to control or *cup-5* RNAi on NGM + 10 mM tryptophan. Data represent mean ± SD. The number of animals analyzed is indicated. ****p < 0.0001 (Student’s t-test). Scale bar: 100 μm.

**Figure S6. Bacterial tryptophan metabolism promotes lipid breakdown in high-fat diet (HFD)-fed mice. Related to Figure 6**.

**(A)** qPCR analysis of total bacterial abundance in fecal samples of HFD-fed mice before (week-4) or after (week-6) antibiotics treatment. Values for each group are normalized to total 16S rRNA levels.

**(B)** Body weight changes in K12- and tnaA-exposed littermates during HFD feeding.

**(C)** Relative liver weight (as a percentage of body weight) in K12- and tnaA-exposed littermates after 12 weeks of HFD feeding. n=4 mice/group.

## Supplemental Table

**Table S1.** RNA-seq data related to Figure 3 and Figure S3.

## STAR★Methods

### Experimental model and study participant details

#### *C. elegans* strains and maintenance

Unless otherwise noted, nematodes were maintained at 20 °C on standard nematode growth medium (NGM) agar plates seeded with *E. coli* OP50 or *E. coli-*K12.

1) The following strain/alleles was obtained from *Caenorhabditis Genetics Center* (CGC): N2; VC4077: lbp-8(gk5151); RB1899: acs-2(ok2457); CF1553: sod-3p::GFP(muIs84); CL2166: gst-4p::GFP::NLS(dvIs19);
2) The following strain/alleles was obtained from Dr. Xiaochen Wang lab and Dr. Chonglin Yang lab: XW5399: *P_ced-1_NUC-1::CHERRY(qxIs257);* XW19180: *P_hs_NUC-1::pHTomato(qxIs750);*
3) The following strains were generated in our lab: YNU623: P*lbp-8*::GFP;P*odr-1*::RFP(*ylfIs46*) YNU624: P*lbp-8*::*lbp-8*::GFP;P*odr-1*::RFP(*ylfIs47*) YNU606: Ex[*acs-2*P::*acs-2*::GFP;*rol-6*](ylfEx325);

### Bacterial strains

Bacterial strains (*E. coli* OP50, *E. coli* K-12 BW25113, Keio collection mutants, and *E. coli* HT115) were grown overnight in LB medium at 37 °C. Cultures were then spread onto standard NGM plates or the specialized agar plates described in this study.

### Cell culture

HUH7 cells were maintained at 37 °C in DMEM (VivaCell C3113-0500) supplemented with 10% fetal bovine serum (FBS; VivaCell C04001-500) and penicillin–streptomycin (VivaCell C3421-0100). Primary hepatocytes, isolated from C57BL/6 mouse livers, were cultured at 37 °C in Williams’ E medium (Thermo Fisher Scientific 12551032) containing 10% FBS and penicillin–streptomycin. Cells were allowed to adhere fully before any experimental treatments.

### Mice

Male C57BL/6J mice (6–8 weeks old, 18–20 g) were obtained from the Yunnan University Animal Center. Animals were group-housed in a pathogen-free facility under a 12 h light/12 h dark cycle with free access to water and food. All procedures were approved by the Institutional Animal Care and Use Committee of Yunnan University.

## Method details

### Generation of transgenic strains

P*lbp-8*::*lbp-8*::GFP reporter: The *lbp-8* promoterand genomic *lbp-8* coding sequence (933bp) were cloned into pPD49.26-GFP. The injection mix contained 10 ng/µl P*lbp-8*::*lbp-8*::GFP and 50 ng/µl *odr-1*p::RFP as a co-injection marker.

P*lbp-8*:: GFP reporter: The *lbp-8* promoter (490bp) was inserted into pPD49.26-GFP. The injection mix comprised 10 ng/µl P*lbp-8*:: GFP and 50 ng/µl *odr-1*p::RFP.

P*acs-2*::*acs-2*::GFP reporter: The *acs-2* promoter (3972 bp) and genomic *acs-2* sequence were cloned into pPD49.26-GFP. The injection mix contained 10 ng/µl P*acs-2*::*acs-2*::GFP and 50 ng/µl rol-6(su1006) as a transformation marker.

### Preparation of worm food with various treatments

Preparation of *E. coli* on NGM plate: standard overnight culture of *E. coli* (wild-type: OP50, K12, or tnaA mutant) grown LB broth was spread onto each NGM plate.

Preparation of *E. coli* on LB plate: standard overnight culture of *E. coli* (wild-type: OP50, K12, or tnaA mutant) grown LB broth was spread onto each LB plate.

Preparation of tryptophan-supplemented NGM plates: Dissolve tryptophan (BBI A601911-0050) into standard nematode growth medium (NGM) to a final concentration of 10 mM, then autoclave. Cool the medium to approximately 55 °C, pour into Petri dishes, and allow the agar to solidify. Once set, seed each plate with an overnight culture of *E. coli* (wild-type: OP50, K12, or tnaA mutant). Leave the plates at room temperature for at least 12 hours to allow the bacterial lawn to establish and dry before use in nematode assays.

Preparation of Heat-killed *E. coli* on NGM plates: Following an established protocol to prepare heat-killed (HK) *E. coli* (Geng et al., 2022). A standard overnight culture of *E. coli* OP50, *E. coli* K12, grown in LB broth was concentrated to 1/10 vol and was then heat-killed at 80°C for 120 min. About 150 ul of the heat-killed bacteria was spread onto each 3.5cm NGM plate with or without tryptophan supplementation.

### Fluorescence intensity measurement

Synchronized L1-stage reporter worms (P*lbp-8*::GFP, P*sod-3*::GFP, or P*acs-2::acs-2::*GFP) were cultured in the indicated media at 20 °C until the L4 stage. For imaging, worms were immobilized in 10 mM levamisole and mounted on agarose pads. Fluorescence was captured under excitation on an Olympus BX53 microscope fitted with a DP80 camera. Whole worm fluorescence intensity was quantified in ImageJ, with at least 30 worms analyzed per reporter strain.

### Quantification of the nucleation rate of LBP-8 protein

Synchronized L1-stage P*lbp-8::lbp-8::*GFP reporter worms were cultured in the specified media at 20 °C until the L4 stage. Individual animals were imaged on a Zeiss LSM 900 laser-scanning confocal microscope. Nuclear and cytoplasmic GFP intensities were quantified using Zeiss ZEN Blue software to calculate the nucleus-to-cytoplasm fluorescence ratio. At least 20 worms were analyzed per condition.

### *E. coli* Keio collection screen

The *E. coli* Keio collection, a library of single-gene knockout mutants (Baba et al., 2006), was used to screen for genes affecting the host’s *lbp-8* expression. For the screen, individual Keio strains were grown overnight at 37 °C in LB medium supplemented with 50 µg/mL kanamycin. A 150 µL aliquot of each culture was spread onto 35 mm LB agar plates. Synchronized L1-stage *C. elegans* carrying the P*lbp-8*::GFP reporter were then transferred onto these plates and incubated at 20 °C for 60 h. Mutants that consistently reduced the P*lbp-8*::GFP expression were selected for follow-up validation assays.

### RNAi treatment in *C. elegans*

All RNAi-by-feeding experiments used *E. coli* HT115 clones from the MRC RNAi Library (Kamath et al., 2003) or the ORF-RNAi Library (Rual et al., 2004). RNAi bacterial cultures were grown overnight at 37 °C in LB medium supplemented with 50 µg/mL carbenicillin, then seeded onto NGM agar plates containing 1 mM IPTG and 50 µg/mL carbenicillin. Synchronized L1-stage worms were transferred onto these RNAi plates and maintained at 20 °C. Worms were allowed to develop either to the L4 stage or the next generation, depending on the specific experimental requirements.

### RNA-seq preparation and analysis

1) Preparation of *C. elegans* samples for RNA-seq

RNA sequencing was conducted with triplicate biological replicates, each representing independent experimental batches.

Group 1 (NGM-based cultivation): Synchronized L1 wild-type animals grown on NGM plate (seeded with *E. coli* K12 or tnaA mutant) supplemented with or without tryptophan.

Group 2 (LB-based cultivation): Synchronized L1 wild-type animals grown on LB plate (seeded with *E. coli* K12 or tnaA mutant) to L4 stage.

All cultures were maintained at 20°C until L4 developmental stage (48–60 h post-synchronization), as determined by vulval morphology. Animals were then flash-frozen in TRIzol® (Invitrogen) for RNA extraction.

2) RNA sequencing and data processing

All sequencing was performed at Biomarker Technologies (BMKGENE). For the RNA sequencing assay, the reference genome index was built using HISAT2 v2.0.5, and paired-end clean reads were aligned to the reference genome with HISAT2 v2.0.5. Differential gene expression analysis was conducted using DESeq2, with p-values adjusted using the Benjamini-Hochberg method (Benjamini and Hochberg, 2018) to control the false discovery rate. Genes with an adjusted p-value ≤ 0.05 were considered differentially expressed. Gene Ontology (GO) enrichment analysis and KEGG pathway enrichment were performed using the clusterProfiler package, with GO terms and KEGG pathways considered significantly enriched if p < 0.05.

### Oil Red O (ORO) staining

#### C. elegans

Young adult worms were collected, washed twice with M9 buffer, and fixed in 0.5% paraformaldehyde (PFA) (BBI, A500684-0500) for 30 minutes following three freeze-thaw cycles in liquid nitrogen. Worms were then washed with M9 to remove residual PFA and incubated with 60% ORO working solution at room temperature in the dark for 30 minutes. The 60% ORO working solution was freshly prepared by diluting ORO stock solution (Sigma, #O1391) with water, followed by rocking and filtration through a 0.22-µm filter. After incubation, worms were washed three times with M9 and mounted onto 2% agarose pads for imaging.

#### Mammalian Cells

Primary hepatocytes, isolated from C57BL/6 mouse livers, were cultured at 37 °C in Williams’ E medium (Thermo Fisher Scientific 12551032) containing 10% FBS and penicillin–streptomycin. Cells were allowed to adhere fully before treatments. The primary mouse hepatocytes were co-treated with palmitic acid (400uM) and supernatants from *E. coli* (K12, or tnaA ) LB cultures (vol of bacterial supernatants: vol of cell cultures=1:10) or only LB medium for 12 hours. Then, cells were washed three times with PBS after removing the culture medium, then fixed with 4% PFA for 1 hour. Following fixation, cells were washed three times with PBS for 5 minutes each and soaked in 60% isopropanol for 2 minutes before incubation with 60% ORO working solution at 37 °C for 15 minutes. After staining, cells were washed three times with PBS, and nuclei were stained with hematoxylin (Servicebio, #G1004). Coverslips were mounted using 80% glycerin before imaging.

#### Mouse Liver Tissue

Liver samples were soaked in 30% sucrose solution overnight, embedded in an optimal cutting temperature (OCT) compound, and sectioned into 10-µm thick slices using a cryostat. Sections were stored at -80 °C until staining.

Before staining, sections were washed with running water to remove OCT and soaked in 60% isopropanol for 5 minutes. Tissue sections were then incubated with ORO working solution at room temperature for 15 minutes, followed by washing with 60% isopropanol until the background of normal liver tissue appeared clear. Sections were counterstained with hematoxylin for 15 seconds and mounted with 80% glycerin before imaging.

Prepared samples were examined using an Olympus BX53 microscope equipped with a DP80 camera. To quantify ORO staining intensity, mean intensity was measured using ImageJ software.

### Detection of oxidative stress in *C. elegans*

MitoTracker™ Red CMXRos is used as an indicator of mitochondrial membrane potential. Under conditions of excessive mitochondrial ROS production, oxidative damage leads to a loss of membrane potential, resulting in reduced dye retention and diminished fluorescence intensity (Nayak et al., 2016).

MitoTracker™ Red CMXRos staining was performed according to a previously published method with some modifications (Liu et al., 2024). A 1 mM stock solution of MitoTracker™ Red CMXRos was prepared in DMSO. To prepare the staining solution, 0.4 µL of the stock solution was added to 50 µL of M9 buffer and mixed thoroughly. Approximately 30–40 worms were then transferred into the solution and incubated in the dark for 5–10 minutes. After staining, the worms were placed on agar plates at 20°C for a 30-minute recovery period. Fluorescence imaging was performed using an Olympus BX53 microscope equipped with a DP80 camera, capturing signals in the RFP fluorescence channel.

### Quantification of lysosomal morphology

NUC-1::mCherry reporter worms at different ages (days 1, 3, and 6) were imaged using laser scanning confocal microscopy (LSM 900, Carl Zeiss). Serial optical sections were analyzed, and the relative area of NUC-1::mCherry-positive vesicular lysosomes and the length of per lysosomal tubules were quantified per unit area 31×43 um^2^ using Zeiss ZEN Blue software.(the relative rare of vesicular lysosome=Sum of vesicular lysosome areas in unit area/unit area 31×43 um^2)^) .The length of tubular lysosomes were randomly quantified at least 10 in each unit area .At least 10 worms were scored in each group at each age.

### Quantification of NUC-1::pHTomato intensity

Day 2 adult *C. elegans* expressing P*hs*NUC-1::pHTomato were incubated at 33°C for 30 minutes, followed by a 24-hour recovery period at 20°C before analysis. Worms were imaged using laser scanning confocal microscopy (LSM 900, Carl Zeiss). The average pHTomato fluorescence intensity per lysosome was quantified using Zeiss ZEN Blue software. At least 20 worms were analyzed for each condition.

### *E. coli* tryptophan metabolite treatment in cell lines

*E. coli*-K12 or *E. coli*-tnaA strains were cultured overnight in LB medium supplemented with or without 50 mM tryptophan. LB medium alone served as the vehicle control. Following overnight incubation, bacterial cultures were centrifuged at 4,000 × g for 15 min to collect supernatants, which were then filter-sterilized through 0.22-µm filter membranes.

For cell treatments, cultures were exposed to the following group:

*E. coli-*K12 culture supernatants derived from LB medium supplemented with tryptophan (Trp-K12-sup); vol of supernatants: vol of cell cultures=1:10.

*E. coli-*tnaA culture supernatants derived from LB medium supplemented with tryptophan (Trp-tnaA-sup); vol of supernatants: vol of cell cultures=1:10.

*E. coli-*K12 culture supernatants from LB medium lacking tryptophan (K12-sup); vol of supernatants: vol of cell cultures=1:10.

Sterile LB medium alone (control); vol of LB: vol of cell cultures=1:10.

### LysoSensor Green staining in cell lines

The fluorescent probe LysoSensor™ Green DND-189 (Thermo Fisher, L7535) is used to measure lysosomal pH. With a pKa of ∼5.2, this dye becomes more fluorescent in acidic environments. To perform staining, the cell culture medium was removed, and cells were washed twice with PBS. The cells were then incubated in 500 µL of DMEM containing 1 µM LysoSensor™ Green DND-189 at 37°C for 1–2 minutes. Fluorescence images were captured using laser scanning confocal microscopy (LSM 900, Carl Zeiss), and fluorescence intensity was quantified using ImageJ software.

### Quantification of NUC-1::CHERRY cleavage

Approximately 200 Day 1 adult worms expressing NUC-1::CHERRY were washed with M9 buffer. Worms were lysed through three cycles of freezing and thawing, followed by boiling with SDS loading buffer. The worm lysate was analyzed by Western blot using anti-CHERRY antibodies (Proteintech, 26765-1-AP, 1:3000) and anti-tubulin antibodies (Sigma, T5168,1:5000). The intensities of NUC-1::CHERRY and free CHERRY bands were quantified using ImageJ software. Cleavage efficiency was calculated as the ratio of CHERRY to the total NUC-1::CHERRY and CHERRY signal.

### Magic Red Staining

Magic Red stain (Abcam, ab270772) is a cell-permeable dye that enters cells and organelles, where it is cleaved by cathepsin B, generating a red fluorescent signal in functional lysosomes. Magic Red stain was prepared in 250x DMSO stock following the manufacturer’s instructions.

For worms staining, dilute the 250× stock solution 1:10 in deionized water (diH₂O) to prepare a 25× staining solution. Then, mix 20 μL of the staining solution with 480 μL of M9 buffer and spread it onto a small plate containing 1mL NGM. Allow the plate to dry under a hood. L4-stage *C. elegans* were transferred onto Magic Red-containing plates and incubated overnight. Fluorescent images were captured using an Olympus BX53 microscope equipped with a DP80 camera, and fluorescence intensity was quantified using ImageJ.

For cell lines staining, the 250x Magic Red stock was diluted 1:10 in diH₂O to prepare a 25× staining solution. Then, 20 µL of the staining solution was further diluted in 480 µL of DMEM (20µL stain+480µL DMEM=500µL) . The cell culture medium was removed, and the staining solution was added to the Huh7 cells (1ml diluted staining solution added into 35mm confolcal dish), followed by incubation at 37°C for 45 minutes. Fluorescence images were captured using laser scanning confocal microscopy (LSM 900, Carl Zeiss), and fluorescence intensity was quantified using ImageJ software.

### Detection of CTSD maturation in mammalian cells

Cathepsin D (CTSD) is a lysosomal aspartic protease belonging to the pepsin superfamily. In human cells, CTSD is initially synthesized as pre-pro-CTSD and undergoes glycosylation in the rough endoplasmic reticulum (ER). The signal peptide is cleaved in the ER, generating pro-CTSD, which is then transported to the Golgi and subsequently to the endosome. Within the lysosome, pro-CTSD is further processed into its mature form (m-CTSD) (Khalkhali-Ellis and Hendrix, 2014).

Lysates from Huh7 cells were prepared using radioimmunoprecipitation assay (RIPA) buffer. Protein samples (10–30 µg) were separated by 10% SDS-PAGE and transferred onto polyvinylidene fluoride (PVDF) membranes. Membranes were blocked and incubated overnight with primary antibodies against Cathepsin D (Abcam, ab75852, 1:3000) and β-Actin (Cell Signaling, 4968S,1:5000). After washing with TBS-Tween and incubation with the appropriate HRP-conjugated secondary antibody anti-Rabbit IgG (ABclonal,AS014, 1:10,000) and goat anti-mouse (Invitrogen, 62-6520, 1:10,000) , protein bands were visualized using enhanced chemiluminescence (ECL, Thermo Scientific). The intensities of mature CTSD and pro-CTSD bands were quantified using ImageJ software.

### Animal experiments and procedures

Male C57BL/6J mice (6–8 weeks old, 18–20 g) were obtained from the Yunnan University Animal Center. Mice were fed a high-fat diet (HFD; Research Diets D12108C; 40 kcal% fat, 1.25% cholesterol) for 14 weeks, then randomly assigned to two groups (n = 6 per group) at week 4: HFD-K12 and HFD-tnaA.

To deplete the gut microbiota, mice received oral gavage (200 ul/mice) of a broad-spectrum antibiotic cocktail (antibiotics mixture containing 0.5 mg/mL of metronidazole /Solarbio, M8060/; 1 mg/mL of vancomycin /Solarbio, V8050; 1 mg/mL of ampicillin /Solarbio, A8180; and 0.5 mg/mL of neomycin sulfate /Solarbio, N8090) every other day for two weeks (from week 4 to week 6). Thereafter, animals were gavaged with either *E. coli* K-12 or the *tnaA* mutant strain (concentrate 40ml of bacterial culture solution to 2ml, then gavage 200ul concentrated bacterial solution-about 2×10_7_ bacteria to each mice) and given free access to a tryptophan-supplemented drinking solution (10mM) for six weeks. Body weight and food intake were recorded weekly.

At week 14, mice were euthanized; blood was collected via retro-orbital bleeding, and liver tissues were excised and weighed. Tissues were either snap-frozen in liquid nitrogen for biochemical assays or fixed in 4% paraformaldehyde for histology.

### Fecal DNA extraction and 16S rRNA qPCR analysis

Analysis 16S rRNA was performed according to a previously published method with some modifications (Chen et al., 2024). Fresh murine feces were collected into sterile 2 mL freezing vials and immediately snap-frozen in liquid nitrogen. For bacterial DNA extraction, the DNA was isolated and purified according to the manufacturer’s protocol using the EIANamp Stool DNA Kit (TIANGEN, DP328-02). Quantitative PCR (qPCR) was performed in triplicates using SYBR Green Master Mix (Thermo Fisher, A25742) on a Real-Time PCR QuantStudio1 system with the accompanying software. The abundance of bacteria was determined using qPCR with universal 16S rRNA gene primers. Standard curves were constructed using the 16S rRNA gene of *E. coli* K12, amplified with conserved 16S rRNA primers. It should be noted that qPCR measures the number of 16S gene copies per sample, not the actual bacterial numbers or colony forming units.

Universal primers: the hypervariable regions V3 and V4 of the bacterial 16S rRNA gene:

V3 (338F 5’-ACTCCTACGGGAGGCAGCA-3’) and V4 (806R 5’-GGACTACHVGGGTWTCTAAT-3’).

### Hematoxylin and Eosin (H&E) staining

Liver specimens were fixed in 4% paraformaldehyde overnight at 4°C, embedded in paraffin, and sectioned into 5 µm thick slices. The sections were stained with hematoxylin and eosin (H&E) and the slides were examined using an Olympus BX53 microscope.

### Microscopy

The *C. elegans* screening was performed using an Olympus MVX10 dissecting microscope. Fluorescence images were captured with an Olympus BX53 microscope equipped with a DP80 camera. Confocal images were acquired using an inverted Zeiss LSM 880/900 confocal microscope system, fitted with an alpha Plan-Apochromat 63× oil immersion objective lens.

### Statistical analysis

All experiments were performed independently at least three times with similar results. Statistical analyses were performed using Student’s t-test or one-way analysis of variance (ANOVA). All analyses were conducted using GraphPad Prism software.

Staining quantification was carried out using ImageJ and ZEN software. Data are presented as mean ± standard error of the mean (SEM). P-values < 0.05 were considered statistically significant. *p<0.05, **p<0.01, ***p<0.001, ****p<0.0001, and “ns” indicates no significant difference.

