## Supplemental Figures for "Microbial Tryptophan Metabolism Activates Host Lysosomal Activity to Facilitate Lipid Breakdown and Ameliorate Hepatic Steatosis"

**Figure S1**

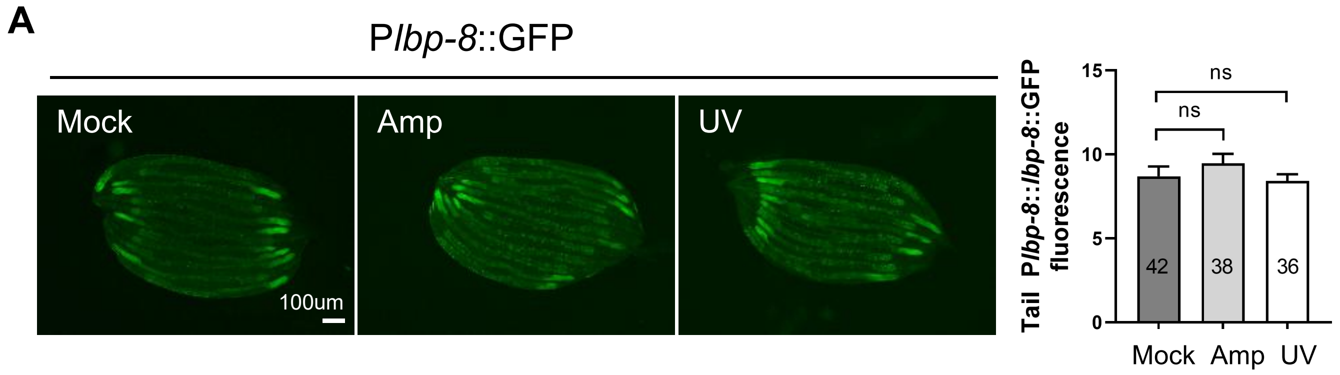


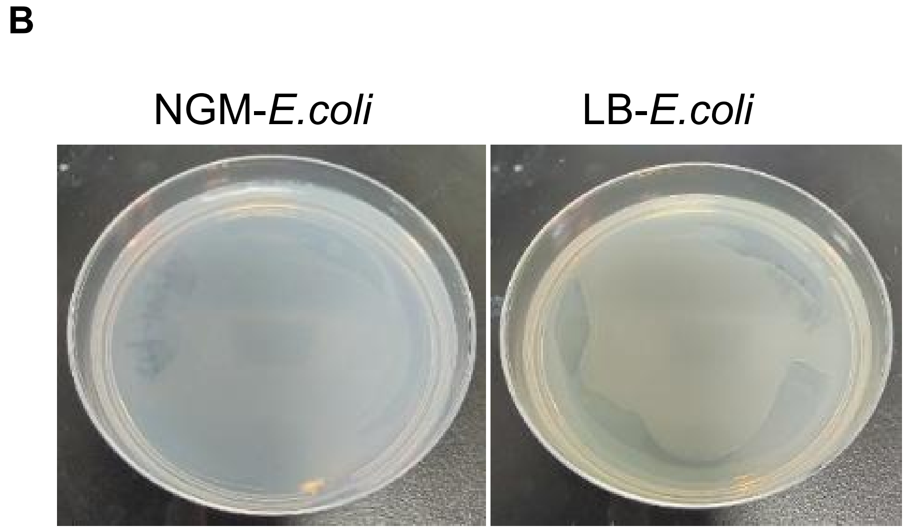


**Figure S1. P*lbp-8::*GFP reporter expression on metabolically compromised *E. coli, and* Bacterial growth state on NGM and LB plate. Related to Figure 1.**

**(A)** Representative fluorescence images of the *Plbp-8::*GFP reporter in worms fed with metabolically compromised *E. coli* (treated with ampicillin or UV-killed). "ns" indicates no significant difference ($p > 0.05$, Student's *t*-test). Data represent mean ± SD. The number of analyzed animals is indicated. Scale bar: 100 μm.

**(B)** Bacterial growth state on NGM and LB culture media seed with *E. coli*-K12.

### Figure S2

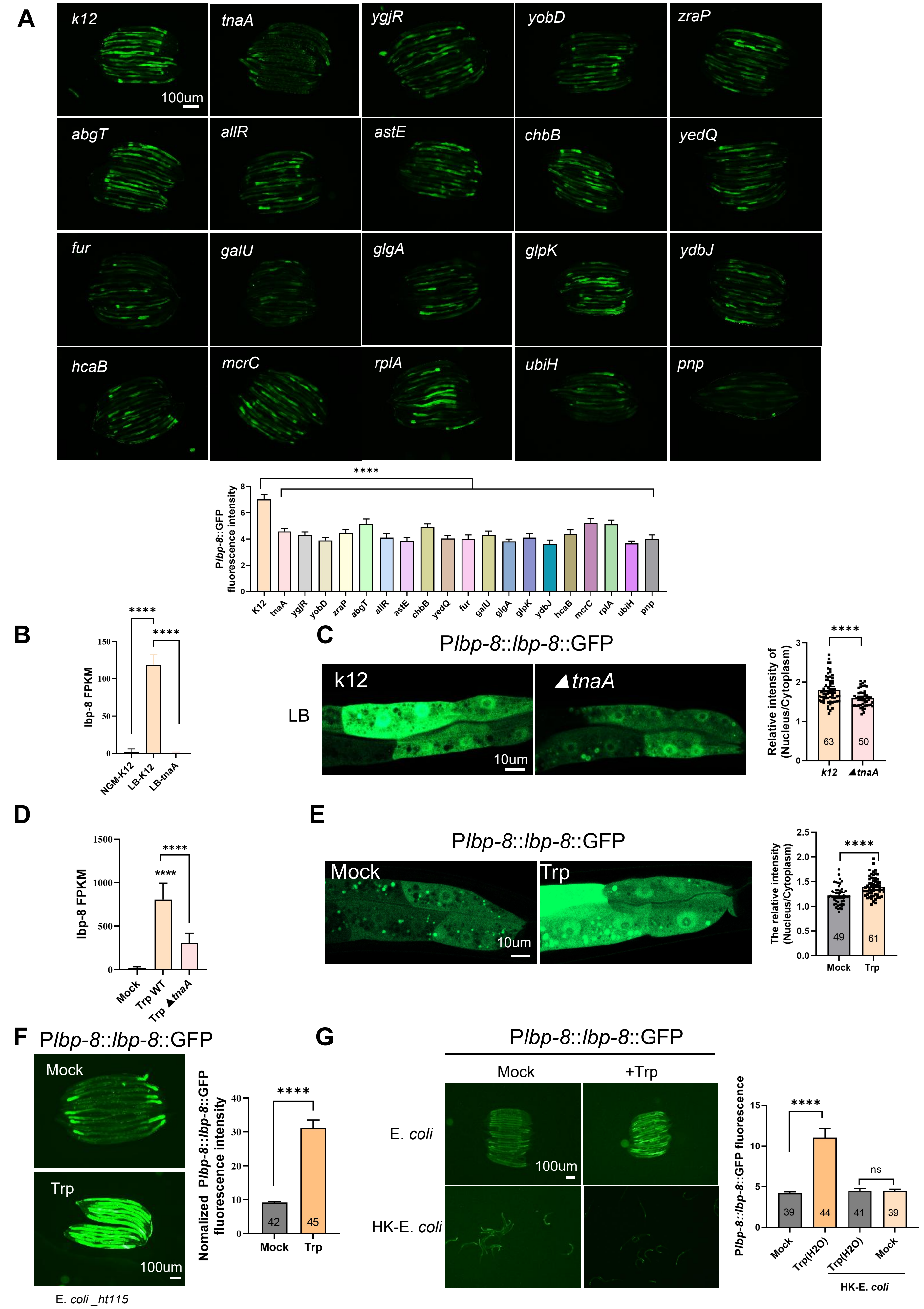

A
k12
tnaA
ygjR
yobD
zraP
100um
abgT
allR
astE
chbB
yedQ
fur
galU
glgA
glpK
ydbJ
hcaB
mcrC
rplA
ubiH
pnp
Plbp-8::GFP fluorescence intensity
****
K12
tnaA
ygjR
yobD
zraP
abgT
allR
astE
chbB
yedQ
fur
galU
glgA
glpK
ydbJ
hcaB
mcrC
rplA
ubiH
pnp
B
lbp-8 FPKM
****
****
NGM-K12
LB-K12
LB-tnaA
C
Plbp-8::lbp-8::GFP
LB
k12
▲tnaA
10um
Relative intensity of (Nucleus/Cytoplasm)
63
50
D
lbp-8 FPKM
****
Mock
Trp WT
Trp ▲tnaA
E
Plbp-8::lbp-8::GFP
Mock
Trp
10um
The relative intensity (Nucleus/Cytoplasm)
49
61
F
Plbp-8::lbp-8::GFP
Mock
Trp
100um
E. coli _ht115
Nomalized Plbp-8::lbp-8::GFP fluorescence intensity
42
45
G
Plbp-8::lbp-8::GFP
Mock
+Trp
E. coli
HK-E. coli
100um
Plbp-8::lbp-8::GFP fluorescence
****
ns
39
44
41
39
Trp(H2O)
HK-E. coli

**Figure S2. Bacterial tryptophan metabolism induce *lbp-8* expression. Related to Figure 2.**

**(A)** Representative fluorescence images and quantification of the P*lbp-8::*GFP reporter in worms fed with *E. coli* K12 mutants, which significantly suppressed *lbp-8* expression on LB medium. Scale bar: 100µm.

**(B)** *lbp-8* mRNA levels in L4-stage wild-type worms exposed to *E. coli* K12 on NGM (NGM-K12) and *E. coli* K12 or *tnaA* cultured on LB medium (LB-K12/LB-*tnaA*), which was extracted from RNA-seq data. Data represent mean ± SD. ****p < 0.0001, Student's *t*-test.

**(C)** Fluorescence images of the P*lbp-8::lbp-8::*GFP reporter in worms exposed to wild-type *E. coli* K12 or *tnaA* mutants on LB medium, along with quantification of nucleus-to-cytoplasm GFP intensity of LBP-8 protein. Data represent mean ± SD. ****p < 0.0001, Student's *t*-test. The number of animals analyzed is indicated. Scale bar: 10 µm.

**(D)** *lbp-8* mRNA levels in L4-stage wild-type worms exposed to *E. coli* K12 grown under standard NGM conditions (Mock) or supplemented with 10 mM tryptophan (*E. coli* K12-Trp, *tnaA*-Trp), which was extracted from RNA-seq data. Data represent mean ± SD. ****p < 0.0001, Student's *t*-test.

**(E)** Fluorescence images of the P*lbp-8::lbp-8::*GFP reporter in worms exposed to wild-type *E. coli* on NGM medium with or without 10 mM tryptophan, along with quantification of nucleus-to-cytoplasm GFP intensity of LBP-8 protein. Data represent mean ± SD. ****p < 0.0001, Student's *t*-test. The number of animals analyzed is indicated. Scale bar: 10 µm.

**(F)** Fluorescence images of the P*lbp-8::lbp-8::*GFP reporter in worms exposed to wild-type *E. coli* on NGM medium with or without 10 mM tryptophan. Data represent mean ± SD. ****p < 0.0001, Student's *t*-test. The number of animals analyzed is indicated. Scale bar: 100 µm.

**(G)** Fluorescence images of the P*lbp-8::lbp-8::*GFP reporter in worms exposed to wild-type *E. coli* or heat-killed *E. coli* cultured in LB medium with or without tryptophan. Data represent mean ± SD. ****p < 0.0001, ns indicates no significant difference (p > 0.05, Student's *t*-test). The number of animals analyzed is indicated. Scale bar: 100 µm.

### Figure S3

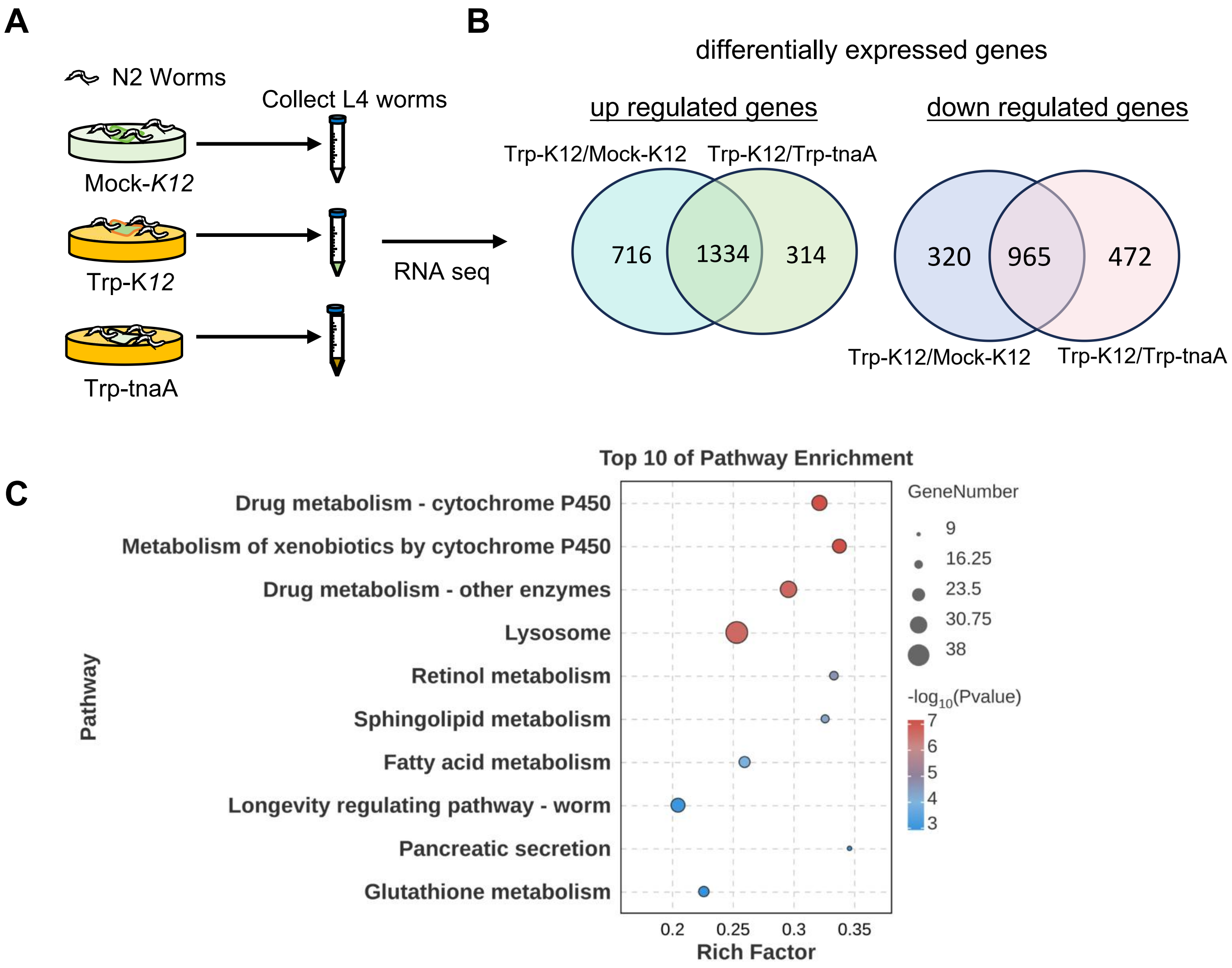


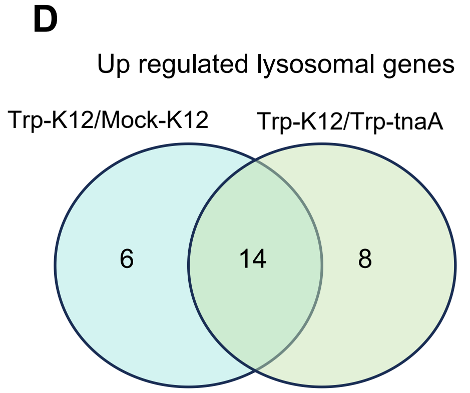


E

| Differential gene | Function in lysosomes |
|---|---|
| *lmp-2* | Intramembrane protease |
| *cpr-3* | Cysteine protease |
| *ppt-1* | Palmitoyl-(protein) hydrolase |
| *lipl-2* | Lipid hydrolase |
| *lipl-6* | Lipid hydrolase |
| *asah-1* | Acylsphingosine amido hydrolase |
| *acp-5* | Acid phosphatase |
| *pho-14* | Acid phosphatase |
| *gba-1* | Glucosylceramidase |
| *gba-2* | Glucosylceramidase |
| *C33G3.4* | Beta-mannosidase |
| *ctsa-1.2* | Serine-type carboxypeptidase |
| *lgmn-1* | LeGuMaiN asparaginyl endopeptidase |
| *T28C12.1* | Transmembrane transporter protein |

F

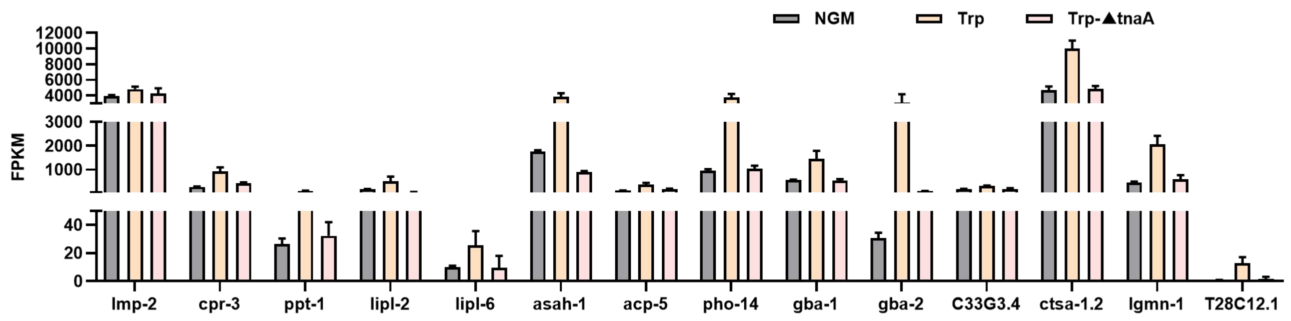

**Figure S3. Bacterial tryptophan metabolism induce lysosomes-related genes expression. Related to Figure 3.**

**(A)** Schematic diagram illustrating the RNA sequencing sample preparation process, including the three experimental conditions: NGM plates seeded with *E. coli* K12 (Mock-K12), tryptophan-supplemented NGM plates seeded with *E. coli* K12 (Trp-K12), and tryptophan-supplemented NGM plates seeded with *E. coli tnaA* (Trp-*tnaA*). Synchronized L1-stage wild-type animals were grown on these plates until reaching the L4 stage, at which point L4-stage animals were collected for sequencing.

**(B)** Venn diagram showing the overlap between genes differentially expressed in Trp-K12 vs NGM-K12 and in Trp-K12 vs Trp-*tnaA*.

**(C)** KEGG enrichment analysis of differentially expressed genes in Trp-K12 vs NGM-K12.

**(D)** Venn diagram displaying the overlap of lysosomal-related genes, which are induced at both group (Trp-K12 vs Mock-K12, Trp-K12 vs Trp-tnaA).

**(E)** List of overlapping lysosomal-related genes.

**(F)** mRNA levels from RNA-seq of the lysosomal-related genes identified in (D).

**Figure S4**

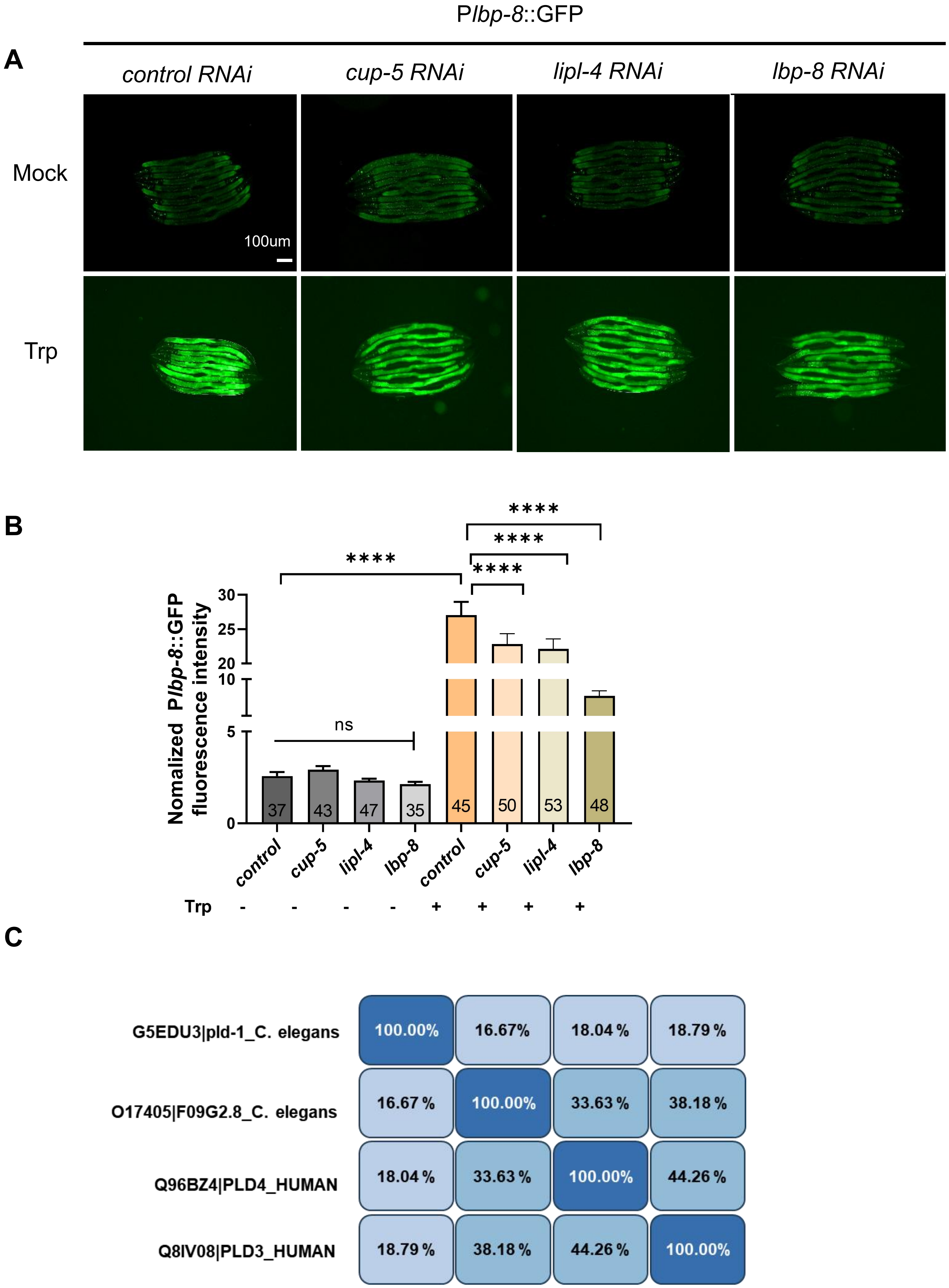

P*lbp-8*::GFP
A
*control RNAi*
*cup-5 RNAi*
*lipl-4 RNAi*
*lbp-8 RNAi*
Mock
Trp
100um
B
Nomalized P*lbp-8*::GFP fluorescence intensity
****
****
****
****
ns
30
25
20
10
5
0
37
43
47
35
45
50
53
48
*control*
*cup-5*
*lipl-4*
*lbp-8*
*control*
*cup-5*
*lipl-4*
*lbp-8*
Trp - - - - + + + +
C
G5EDU3|pld-1_C. elegans
O17405|F09G2.8_C. elegans
Q96BZ4|PLD4_HUMAN
Q8IV08|PLD3_HUMAN
100.00% 16.67% 18.04 % 18.79 %
16.67 % 100.00% 33.63 % 38.18 %
18.04 % 33.63 % 100.00% 44.26 %
18.79 % 38.18 % 44.26 % 100.00%

**Figure S4. Bacterial tryptophan metabolism promotes lipid degradation in a lysosomal-dependent manner. Related to Figure 4.**

**(A-B)** Representative fluorescence images (A) and quantification (B) of the P*lbp-8*::GFP reporter in wild-type L4 worms treated with control, *cup-5*, *lipl-4* or *lbp-8* (negative control) RNAi, grown on NGM plates with or without 10 mM tryptophan. Data represent mean $\pm$ SD. The number of animals analyzed is indicated. ****$p < 0.0001$, "ns" $p > 0.05$ (Student's t-test). Scale bar, 100 μm.

**(C)** Homology alignment of *C. elegans* PLD-1 and F09G2.8 with human PLD-3 and PLD-4 protein sequences. For each row/column, 100% represents the target protein, and percentages indicate sequence similarity relative to the target protein.

### Figure S5

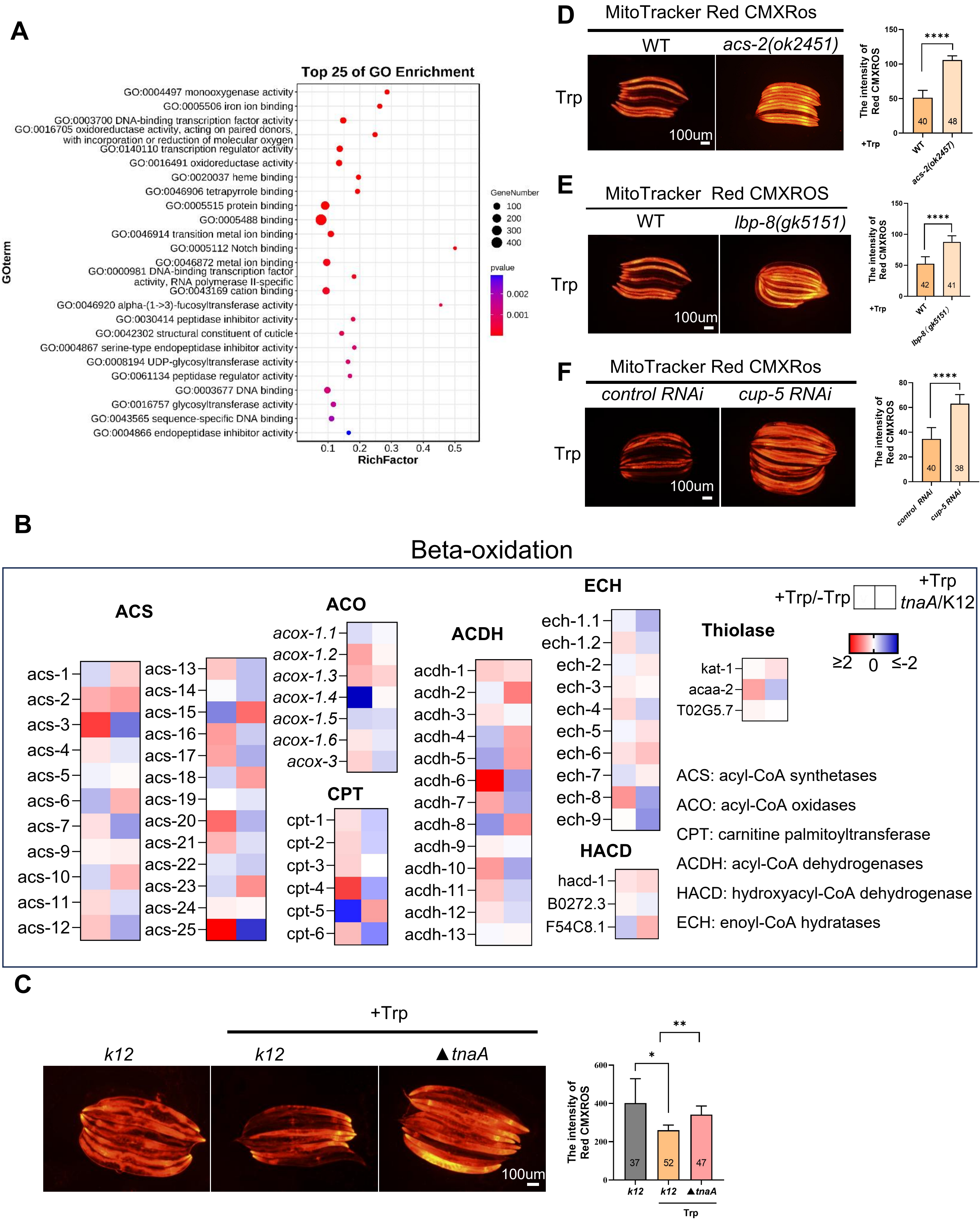

A
Top 25 of GO Enrichment
GO:0004497 monooxygenase activity
GO:0005506 iron ion binding
GO:0003700 DNA-binding transcription factor activity
GO:0016705 oxidoreductase activity, acting on paired donors, with incorporation or reduction of molecular oxygen
GO:0140110 transcription regulator activity
GO:0016491 oxidoreductase activity
GO:0020037 heme binding
GO:0046906 tetrapyrrole binding
GO:0005515 protein binding
GO:0005488 binding
GO:0046914 transition metal ion binding
GO:0005112 Notch binding
GO:0046872 metal ion binding
GO:0000981 DNA-binding transcription factor activity, RNA polymerase II-specific
GO:0043169 cation binding
GO:0046920 alpha-(1->3)-fucosyltransferase activity
GO:0030414 peptidase inhibitor activity
GO:0042302 structural constituent of cuticle
GO:0004867 serine-type endopeptidase inhibitor activity
GO:0008194 UDP-glycosyltransferase activity
GO:0061134 peptidase regulator activity
GO:0003677 DNA binding
GO:0016757 glycosyltransferase activity
GO:0043565 sequence-specific DNA binding
GO:0004866 endopeptidase inhibitor activity
GOterm
RichFactor
GeneNumber
100
200
300
400
pvalue
0.002
0.001
D
MitoTracker Red CMXRos
WT
acs-2(ok2451)
Trp
100um
The intensity of Red CMXROS
****
40
48
+Trp
WT
acs-2(ok2457)
E
MitoTracker Red CMXROS
WT
lbp-8(gk5151)
Trp
100um
****
42
41
lbp-8(gk5151)
F
MitoTracker Red CMXRos
control RNAi
cup-5 RNAi
Trp
100um
****
40
38
B
Beta-oxidation
ACS
ACO
ACDH
ECH
Thiolase
HACD
CPT
+Trp/-Trp
+Trp tnaA/K12
≥2 0 ≤-2
acs-1 acs-2 acs-3 acs-4 acs-5 acs-6 acs-7 acs-9 acs-10 acs-11 acs-12
acs-13 acs-14 acs-15 acs-16 acs-17 acs-18 acs-19 acs-20 acs-21 acs-22 acs-23 acs-24 acs-25
acox-1.1 acox-1.2 acox-1.3 acox-1.4 acox-1.5 acox-1.6 acox-3
cpt-1 cpt-2 cpt-3 cpt-4 cpt-5 cpt-6
acdh-1 acdh-2 acdh-3 acdh-4 acdh-5 acdh-6 acdh-7 acdh-8 acdh-9 acdh-10 acdh-11 acdh-12 acdh-13
ech-1.1 ech-1.2 ech-2 ech-3 ech-4 ech-5 ech-6 ech-7 ech-8 ech-9
hacd-1 B0272.3 F54C8.1
kat-1 acaa-2 T02G5.7
ACS: acyl-CoA synthetases
ACO: acyl-CoA oxidases
CPT: carnitine palmitoyltransferase
ACDH: acyl-CoA dehydrogenases
HACD: hydroxyacyl-CoA dehydrogenase
ECH: enoyl-CoA hydratases
C
+Trp
k12
k12
▲tnaA
100um
The intensity of Red CMXROS
*
**
37
52
47
Trp

**Figure S5. Bacterial tryptophan metabolism enhances mitochondrial β-oxidation via lysosomal activation to facilitate lipid metabolism. Related to Figure 5.**

**(A)** Gene ontology (GO) enrichment analysis of 1334 genes being significantly induced at both condition (Trp-K12 vs Mock-K12, Trp-K12 vs Trp-tnaA) (Figure S3B).

**(B)** Heatmap showing fold changes in mRNA levels of β-oxidation-related genes in worms fed Trp-supplemented K12 (Trp-K12) versus mock-treated K12 (Mock-K12), and Trp-supplemented Δ*tnaA* (Trp-tnaA) versus Trp-supplemented K12 (Trp-K12). Fold changes were calculated by dividing each gene's expression level under the indicated conditions.

**(C)** Representative fluorescence images and quantification of MitoTracker™ Red CMXRos in L4 worms fed on NGM plates seeded with wild-type K12 (control) or with K12/Δ*tnaA* plus 10 mM tryptophan. Data represent mean ± SD. The number of animals analyzed is indicated. $^{**}p < 0.01$, $^{*}p < 0.05$ (Student's t-test). Scale bar: 100 μm.

**(D-E)** Representative microscope images and quantification of MitoTracker™ Red CMXRos fluorescence in wild-type, *acs-2(ok2457)* (D) or *lbp-8(gk5151)* (E) mutants L4 worms fed NGM + 10 mM tryptophan, seeded with *E. coli*-K12. Data represent mean ± SD. The number of animals analyzed is indicated. $^{****}p < 0.0001$ (Student's t-test). Scale bar: 100 μm.

**(F)** Representative microscope images and quantification of MitoTracker™ Red CMXRos fluorescence in wild-type worms subjected to control or *cup-5* RNAi on NGM + 10 mM tryptophan. Data represent mean ± SD. The number of animals analyzed is indicated. $^{****}p < 0.0001$ (Student's t-test). Scale bar: 100 μm.

#### Figure S6

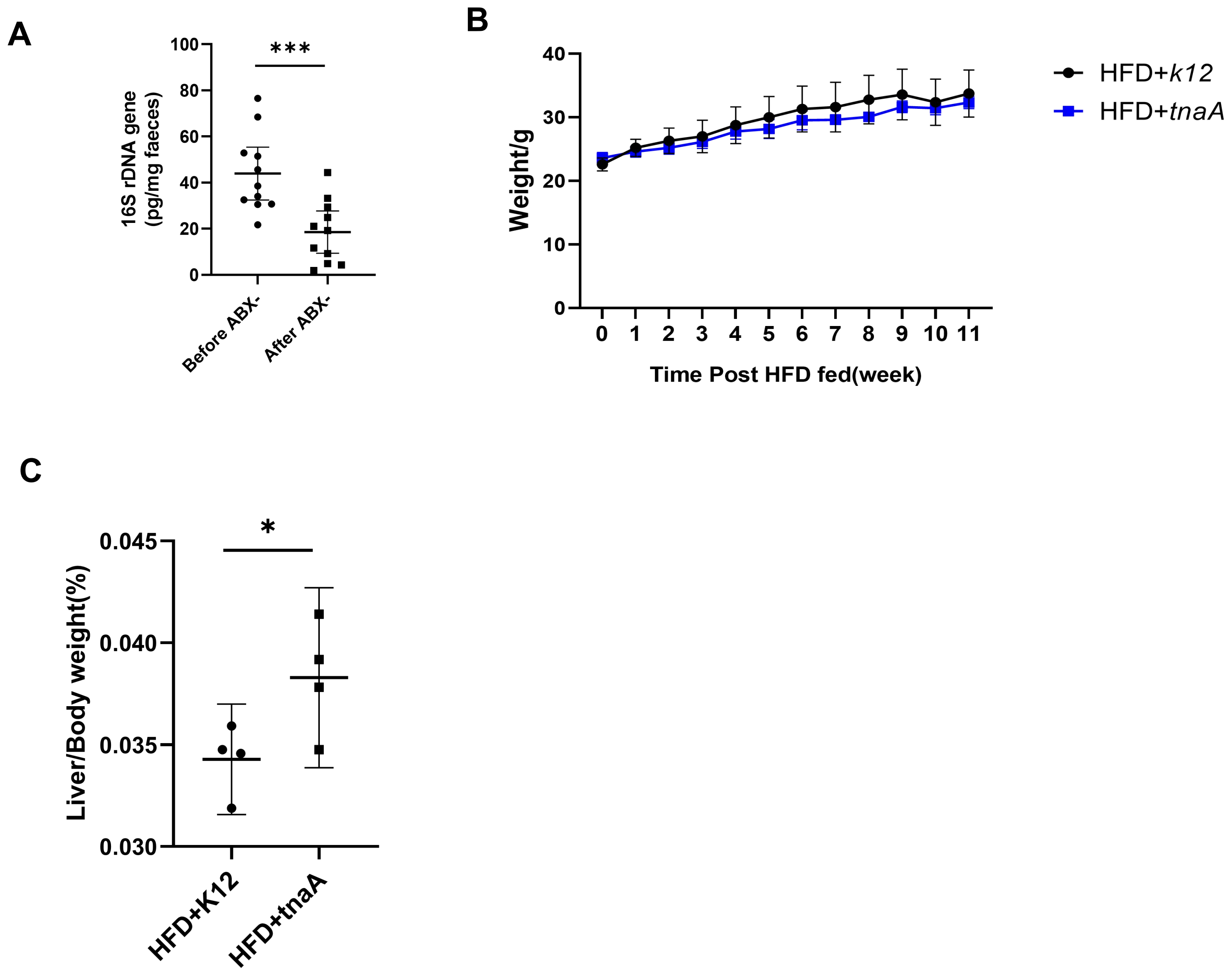


**Figure S6. Bacterial tryptophan metabolism promotes lipid breakdown in high-fat diet (HFD)-fed mice. Related to Figure 6.**

**(A)** qPCR analysis of total bacterial abundance in fecal samples of HFD-fed mice before (week-4) or after (week-6) antibiotics treatment. Values for each group are normalized to total 16S rRNA levels.

**(B)** Body weight changes in K12- and tnaA-exposed littermates during HFD feeding.

**(C)** Relative liver weight (as a percentage of body weight) in K12- and tnaA-exposed littermates after 12 weeks of HFD feeding. n=4 mice/group.
